# Disrupted structure and aberrant function of CHIP mediates the loss of motor and cognitive function in preclinical models of cerebellar CHIPopathy

**DOI:** 10.1101/283192

**Authors:** Chang-he Shi, Carrie Rubel, Sarah E. Soss, Rebekah Sanchez-Hodge, Shuo Zhang, Donte A. Stevens, Holly McDonough, Richard C. Page, Walter J. Chazin, Cam Patterson, Cheng-yuan Mao, Monte S. Willis, Hai-Yang Luo, Mi-bo Tang, Pan Du, Yao-he Wang, Zheng-wei Hu, Yu-ming Xu, Jonathan C. Schisler

## Abstract

CHIP (carboxyl terminus of heat shock 70-interacting protein) has long been recognized as an active member of the cellular protein quality control system given the ability of CHIP to function as both a co-chaperone and ubiquitin ligase. Mutations in CHIP are the driver of spinocerebellar autosomal recessive 16 (SCAR16), or cerebellar CHIPopathy, as we initially discovered this disease was caused by a loss of CHIP ubiquitin ligase function. The initial mutation describing SCAR16 was a missense mutation in the ubiquitin ligase domain of CHIP (p.T246M). Using multiple biophysical and cellular approaches, we demonstrate that T246M mutation results in structural disorganization and misfolding of the CHIP U-box domain, promoting oligomerization, and increased proteasome-dependent turnover. CHIP-T246M has no ligase activity, but maintains interactions with chaperones and alters the co-chaperone function of CHIP. To establish preclinical models of SCAR16, we engineered T246M at the endogenous locus in both mice and rats. Animals homozygous for T246M had both cognitive and motor cerebellar dysfunction distinct from those observed in the CHIP null animal model, as well as deficits in learning and memory, reflective of the cognitive deficits reported in SCAR16 patients. We conclude that the T246M mutation is not equivalent to the total loss of CHIP, supporting the concept that disease-causing CHIP mutations have different biophysical and functional repercussions on CHIP function that may directly correlate to the spectrum of clinical phenotypes observed in SCAR16 patients. Our findings both further expand our basic understanding of CHIP biology and provide meaningful mechanistic insight underlying the molecular drivers of SCAR16 disease pathology, which may be used to inform the development of novel therapeutics for this devastating disease.

## Introduction

Protein quality control (PCQ) involves a specialized cellular surveillance system that monitors protein integrity, identifies unfolded or damaged proteins, and then either repairs or targets them for degradation. CHIP is abundantly expressed in most tissues and plays a central role in maintaining protein quality control [1]. CHIP is uniquely suited as a regulator of protein quality control due to its dual functions as both a co-chaperone protein and ubiquitin ligase enzyme. As a co-chaperone, CHIP interacts with heat shock protein (HSP)-bound proteins to aid in substrate stabilization and refolding [2]. Conversely, as a ubiquitin ligase CHIP ubiquitinates terminally-defective proteins and prepares them for degradation by the Ubiquitin Proteasome System (UPS) [3]. Since the discovery of CHIP in 1999 [1], numerous reports detailing CHIP’s co-chaperone and ubiquitin ligase activities in both the brain and heart have been published [4–7]. However, recent reports describing surprising new roles for CHIP have emerged. These roles include autonomous chaperone activity [8, 9], the regulation of cardiac metabolic homeostasis via the metabolic sensor AMPK (AMP-activated protein kinase) [9], and DNA damage repair [10]. Most recently, CHIP was implicated in the pathophysiology associated with cerebellar CHIPopathy, or spinocerelbellar autosomal recessive 16 (SCAR16) [11, 12], representing the first direct association between a CHIP polymorphism and a human disease.

Cerebellar CHIPopathy (MIM 615768) is a form of autosomal recessive spinocerebellar ataxia that can also be accompanied with hypogonadism, similar to the clinical phenotype of Gordon Holmes Syndrome (GHS) [13]. Despite over 100 years of clinical recognition, only recently have causal mutations for GHS been identified. These mutations include the ubiquitin ligase RNF216 and deubiquitinase OTUD4 [14], which suggest that faulty ubiquitination plays an essential role in the pathophysiology of ataxia. Using exome sequencing, we identified a mutation in STUB1 (the gene encoding CHIP) in two patients initially diagnosed with GHS [11]. We found that this STUB1 mutation (p.T246M) resulted in a loss in the ubiquitin ligase function of CHIP [11]. Combined with our studies demonstrating that mice lacking the expression of CHIP display motor deficiencies and some aspects of the hypogonadism observed in patients with STUB1 mutations, our CHIP knockout mouse represented the first animal model of CHIPopathy [11]. Subsequently, numerous clinical studies identified STUB1 mutations, confirming our initial identification of a new disease [12, 15–22]. Additional studies using in vitro models suggest that CHIP mutations can impart differential biochemical characteristics [23, 24], although how these characteristics relate to cellular function and diseases processes are not known.

The amino acid substitutions reported in cases of cerebellar CHIPopathy result in nonsense, missense, frameshift, or splicing mutations; the majority of which are predicted to significantly alter protein function [12]. Given the clinical heterogeneity of neuroendocrine phenotypes in STUB1 patients, specific CHIP mutations likely have varying biophysical and functional consequences to CHIP function. In this context, an animal model with a total loss of CHIP may not adequately represent the spectrum of human disease represented by SCAR16. For these reasons, we complemented the biophysical and cellular repercussions of CHIP-T246M with two rodent models engineered with CRISPR/Cas9 to mimic this human mutation. Additionally, we performed in-depth behavioral assessments to determine the effects of T246M mutation at a whole-animal level to establish a suitable preclinical model for SCAR16. Studying CHIP mutations both in vitro and in vivo allows us to delineate the contribution of co-chaperone, ubiquitin ligase, and other emerging CHIP activities to specific deficits observed in a disease-relevant context in vivo. These results provide insights that are valuable for the development of effective therapies for this devastating degenerative disease.

## Results

### BIOCHEMICAL AND CELLULAR ANALYSIS OF CHIP-T246M

#### The T246M mutation destabilizes the structure of the U-box and CHIP-T246M forms decamers and dodecamers in vitro and in cells

Asymmetric homodimerization of CHIP as well as conformational flexibility are required for CHIP ubiquitin ligase activity. Critical to both the dimerization and conformational flexibility is the U-box domain [3, 25] where T246 is located and is a highly conserved residue across CHIP homologs [11]. Furthermore, T246 is located in the core of a conserved beta hairpin turn that lies at the interface between the two molecules of the CHIP dimer (Figure 1A). Modeling of the T246M amino acid substitution predicts that this mutation would impact the tertiary structure of the U-box and hence disrupt the formation of functional dimers, consequently reducing or abolishing CHIP’s ubiquitin ligase function towards both chaperone and non-chaperone substrates. To test the effects of T246M substitution on U-box structure, we performed solution NMR on purified WT and T246M CHIP U-box domains. Whereas WT CHIP U-box exhibit distinct peaks spread throughout the 2D ^15^N-^1^H HSQC spectrum characteristic of a stable, structured domain (Figure 1B), the T246M spectrum is collapsed and has broad resonances, suggesting the loss of stable globular structure, multiple conformations and aggregation (Figure 1B). Additionally, circular dichroism spectra were acquired for the isolated WT and T246M CHIP U-box domains. Compared to the secondary structure content of the WT U-box, the signal from T246M U-box was consistent with loss or regular secondary structure and a shift to random coil conformation (Figure 1C). Further, we monitored the protein melting temperature (T_m_) at 222 nm, the characteristic wavelength for α-helices. The WT U-box exhibited a sigmoidal curve consistent with unfolding of a globular, folded domain with a Tm of 30 °C, whereas T246M appeared to be unfolded at room temperature and did not undergo any change with temperature (Figure 1C). Together these data suggest the T246M mutation destabilizes the U-box domain, resulting in a loss of secondary and tertiary structure.

**Figure 1:**
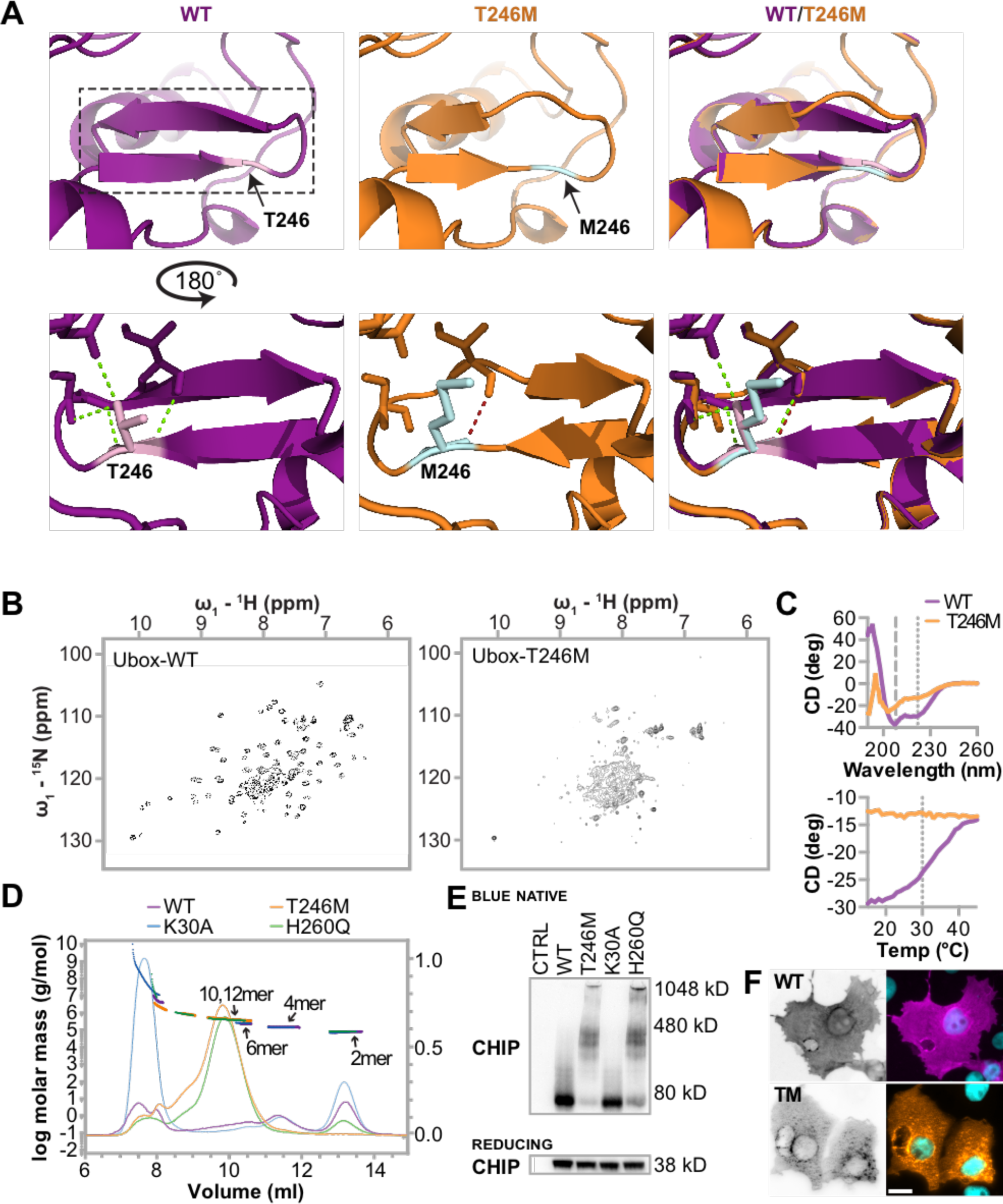
CHIP-T246M disrupts the structure of the U-box and promotes the formation of soluble oligomers. (**A, upper**) The structure of the beta hairpin region that harbors residue 246 is shown in CHIP-WT (purple, **left**) compared to the modeling of CHIP-T246M (orange, **center**). The overlay is also provided (**right**). (**A, lower**) The structures are rotated 180 degrees and the hydrogen bonds that are predicted to be impacted in this region by the T246M mutation are indicated (green). (**B**) 600-MHz ^15^N-^1^H transverse relaxation-optimized spectroscopy-HSQC spectra collected at 293K for ^2^H,^15^N-labeled WT (left) and T246M (right) CHIP U box (218-303). (**C, upper**) Circular dichroism spectroscopy data collected for the U-box of WT (purple) and T246M (orange). (**C, lower**) Melting point determinations for the U-box of WT and T246M CHIP at 222nm. (**D**) Size distribution of full-length proteins of either WT (purple), T246M (orange), K30A (blue) and H260Q (green) CHIP determined by size-exclusion chromatography and multi-angle lights scattering. The molecular mass of oligomeric species of each protein are indicated (left Y-axis, thick lines) and the Rayleigh ratio chromatographs represent the amount of light scattering (right Y-axis, thin lines). (**E**) COS-7 cells were co-transfected with the indicated vectors (CTRL = pcDNA3, WT = pcDNA3-mycCHIP, T246M = pcDNA3-myc CHIP-T246M, K30A=pcDNA3-mycCHIP-K30A, H260Q = pcDNA3-mycCHIP-H260Q). Cells were collected on ice and total protein collected and freshly separated by BN PAGE under native conditions or SDS-PAGE under reducing conditions and immunoblotted with an anti-myc (CHIP). Locations of molecular weight standards in kilodaltons (kD) are indicated. (**F**) COS-7 cells were co-transfected with the indicated transgenes. 24 hours post transfection cells were fixed and immunostained for myc-CHIP expression (scale bar =20 microns).

Previous size exclusion chromatography (SEC) analysis suggested that full-length CHIP harboring the T246M formed large aggregates greater than 670 kDa [23, 24], however, SEC alone cannot account for differences in protein conformation or account for protein-column interactions [26]. Given the known dynamics of CHIP conformation in solution, we used an analytical approach, SEC coupled with multi-angle light scattering, to determine true molar mass and radius in solution and compared these data to the composition of native CHIP-T246M when expressed in cells using blue native PAGE and immunoblot analysis. As expected, both the WT protein and the K30A control mutant were predominantly dimers, 72 kDa (Figure 1D, Table 1). However, the U-box domain mutants T246M and H260Q were detected as higher-order oligomers, suggesting CHIP-T246M exist in cells predominantly as 10-12mers (Figure 1D). Native gels of WT CHIP and the same point mutants expressed in COS7 cells also confirmed that CHIP-WT and CHIP-K30A form dimers whereas CHIP-T246M and CHIP-H260Q form higher-order oligomers (Figure 1E). Next, we performed indirect immunofluorescence to observe CHIP localization and expression patterns in the same cell model. Not surprisingly, CHIP-WT protein is detected as diffuse staining throughout the cytoplasm and within the nucleus, while CHIP-T246M protein appears as punctate staining in the cytoplasm and perinuclear regions, perhaps reflective of CHIP-T246M oligomers (Figure 1E). Taken together, these data suggest that the mutation destabilizes the U-box domain resulting in the loss of ubiquitin ligase activity of T246M CHIP previously observed [11] and promotion of oligomerization.

**Table 1:** SEC-MALS analysis of CHIP proteins. The true molecular weight (mw) in Daltons is provided along with the fractional contribution of each oligomeric species (Xmer) of CHIP identified. With some proteins, a small amount was detected as very high molecular weight (VWMW).

| Protein | 2mer | 4mer | 6mer | 10mer | 20mer | 50mer | VMMW |
| --- | --- | --- | --- | --- | --- | --- | --- |
| WT (mw) | 7.62E+04 | 1.43E+05 | 2.32E+05 |  |  |  | 4.87E+06 |
| WT (%) | 69.7% | 25.2% | 4.8% |  |  |  | 0.4% |
| K30A (mw) | 7.31E+04 | 1.38E+05 | 2.51E+05 |  |  |  | 1.86E+07 |
| K30A (%) | 80.4% | 16.4% | 2.6% |  |  |  | 0.7% |
| T246M (mw) |  |  |  | 4.13E+05 | 8.01E+05 | 1.99E+06 |  |
| T246M (%) |  |  |  | 91.0% | 7.4% | 1.6% |  |
| H260Q (mw) | 8.13E+04 |  |  | 3.97E+05 | 8.29E+05 |  |  |
| H260Q (%) | 28.5% |  |  | 67.8% | 3.5% |  |  |

#### The T246M mutation increases co-chaperone activity

When cells are exposed to heat, the co-chaperone activity of CHIP initiates the heat shock response through the activation of heat shock factor 1 (HSF1). At the same time, heat induces the oligomerization of CHIP both in vitro and in cells, enhancing the chaperone activity of the protein while not affecting ubiquitin ligase activity [8]. Remarkably, the oligomerization of CHIP-WT that occurs with heat resembles the oligomerization of CHIP-T246M and –H260Q [8]. Given T246M increased the interaction between CHIP and HSC70/HSP70 [11] and the oligomeric status of CHIP-T246M (Figure 1E), we hypothesized that CHIP-T246M would still have the ability to co-chaperone HSF1. To test whether CHIP’s regulation of HSF1 remains intact with T246M, we transiently expressed CHIP constructs in COS7 cells and measured the nuclear translocation of HSF1 and changes in HSF1 transcriptional activity. As previously reported [27] we observed that expression of WT CHIP promotes the nuclear translocation of HSF1 (Figure 2A) and stimulates HSF1 activity (Figure 2B). Remarkably, expression of CHIP-T246M also promoted HSF1 nuclear translocation (Figure 2A) and enhanced HSF1 transcriptional activity (Figure 2B). In contrast, abolishing chaperone interactions via the K30A mutation of CHIP did not affect HSF1 dynamics. The ability of T246M to affect HSF1 was dependent on a functional TPR domain, as the effect of T246M on HSF1 activity is lost with the K30A-T246M double mutant (Figure 2B).

**Figure 2:**
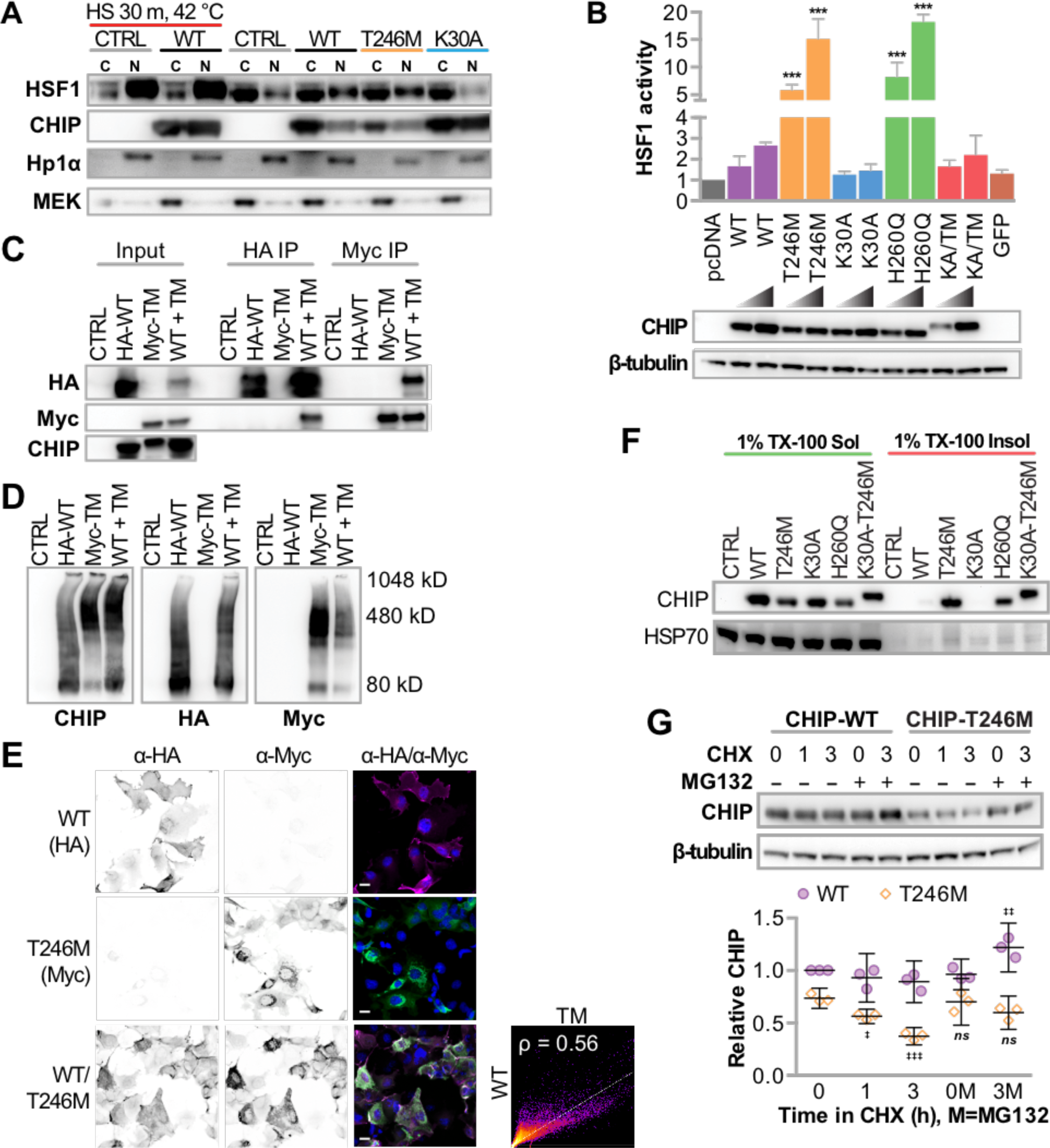
Cellular characterization of CHIP-T246M revealed enhanced activation of HSF1, changes in solubility, and increased turnover. (**A**) Immunoblot analysis of HSF1, CHIP, HP1α (nuclear marker) and MEK (cytosolic marker) in cytosolic (C) and nuclear (N) fractions from COS-7 cells transiently transfected with the indicated vectors (CTRL = pcDNA3, WT = pcDNA3-mycCHIP, T246M = pcDNA3-mycCHIP-T246M, K30A = pcDNA3-mycCHIP-K30A) treated with or without heat shock as indicated. (**B**) Bar graph of HSF1 transcription activity represented by the mean ± 95% CI normalized to control vector (pcDNA) conditions in COS-7 cells transiently transfected with increasing amount if DNA using the indicated vectors (K30A-T246M = pcDNA3-mycCHIP-K30A-T246M, GFP = green fluorescent protein), N = 4 biological replicates, *** *p* < 0.0001 via Dunnet’s multiple comparison test to WT conditions. Immunoblot analysis of the myc-tag and β-tubulin confirmed transgene expression. (**C**) COS-7 cells were co-transfected with the indicated vectors (transgenes, HA-WT = pcDNA3-HA-CHIP, Myc-TM = pcDNA3-mycCHIP-T246M, WT + TM = HA-WT and Myc-TM). CHIP protein was immunoprecipitated with Anti-HA or Anti-Myc affinity gel. The inputs and resulting precipitants (IP) were separated by SDS-PAGE and immunoblotted with the indicated antibodies. (**D**) COS-7 cells were co-transfected with the indicated transgenes and cell lysates were separated by BN-PAGE and immunoblotted with the indicated antibodies. Locations of molecular weight standards in kilodaltons (kD) are indicated. (**E**) Micrographs of indirect immunofluorescence from COS-7 expressing the indicated transgenes to detect CHIP-WT (left), CHIP-T246M (center), or both (right). DAPI nuclear staining is also included (right panels) and the scale bar represents 20 microns. Co-localization of HA-WT and Myc-TM is represented by scatter plot and the indicated Pearson correlation (ρ). (**F**) Solubility analysis (in 1% Triton X-100, TX-100) of CHIP proteins, HSP70, and AMPKα in COS-7 cells determined by immunoblot analysis. (**G**) Immunoblot analysis of CHIP expression (**top**) in COS-7 cells treated with 50 µg/ml cycloheximide (CHX) for the indicated time (h) in the presence or absence of 20 µM MG132. β-tubulin. Densitometry analysis (**bottom**) represented by dot plot and summarized by the mean ± 95% CI of relative levels of CHIP protein were normalized to β-tubulin, N = 3 biological replicates, ‡, ‡‡, ‡‡‡ correspond to *p* < 0.05, 0.01, 0.0001 compared to starting expression levels within each construct via Tukey’s post hoc test.

#### Co-expression of CHIP-WT and CHIP-T246M does not alter oligomerization or localization

The recessive basis of SCAR16 suggests that the wild-type allele is sufficient to overcome the disease-bearing mutated allele. Given the U-box is important to dimerization of CHIP we hypothesized that CHIP-WT would not interact with CHIP-T246M. Interestingly, using extracts from cells expressing both CHIP-WT and CHIP-T246M, these two proteins co-immunoprecipitated each other (Figure 2C) suggesting the two forms of CHIP can interact with each other. However, when we directly separated the lysates via BN-PAGE immunoblot analysis, we observed only subtle differences in oligomerization patterns when CHIP-WT and CHIP-T246M are co-expressed compared to the single protein expression conditions (Figure 2D). Given that BN-PAGE can limit antibody detection of proteins given the native conditions, possibly obscuring changes in visualizing heterocomplexes, we also analyzed expression via indirect immunofluorescence, and we observed that the localization of CHIP-WT was unaffected by co-expression of CHIP-T246M (Figure 2E) despite extensive overlap in the localization of CHIP expression within cells (Figure 2E). Together, these data suggest that while WT and T246M CHIP interact in crude extracts, the localization and dimerization status of CHIP-WT is largely unaffected by the presence of CHIP-T246M. Therefore, CHIP function likely also remains intact in the heterozygous condition and the presence of WT CHIP does not appear to alter the oligomerization or distribution of CHIP-T246M, consistent with the recessive nature of this disease-causing mutation.

#### Change in solubility and increased proteasome-dependent turnover of CHIP-T246

We consistently observed lower levels of soluble CHIP-T246M, CHIP-H260Q, and CHIP-K30A/T246M protein relative to CHIP-WT when transiently transfecting equal amounts of vector DNA (Figure 2B). We hypothesized that this decrease in soluble CHIP-T246M expression could be due to changes in solubility and/or stability. To test for changes in solubility, we performed SDS-PAGE and immunoblot analysis of CHIP in both the soluble and insoluble fraction of whole cell lysates. We observed CHIP-T246M, -H260Q, and -K30A-T246M CHIP in both the soluble and insoluble fraction; in contrast, CHIP-WT or -K30A was found only in the soluble fraction (Figure 2F). In addition to the change in solubility, we also observed a shorter half-life of soluble CHIP-T246M compared to CHIP-WT, an effect that that was dependent on proteasome activity (Figure 2G). Taken together, our data suggest that the T246M mutation results in altered cellular distribution, solubility, and stability and likely contributes to the loss of CHIP function.

#### Protein levels of CHIP-T246M are reduced in fibroblasts from SCAR16 patients and mice engineered with the equivalent mutation

Prior to this report, analyses of CHIP-T246M by others were limited to in vitro analyses or exogenous expression of CHIP in cells that robustly express endogenous CHIP [23, 24]. Thus, we utilized primary embryonic fibroblasts (MEFs) isolated from mice engineered with the corresponding murine amino acid substitution at the endogenous CHIP locus (T247M) [28]. Consistent with our exogenous expression models, soluble CHIP protein levels were robustly reduced in MEFs isolated from different M246/M246 mouse embryos relative to either T246/T246 or T246/M246 littermates (Figure 3A). The decrease in steady-state CHIP in M246/M246 MEFs appeared to be at the post-translational level, as mRNA levels of *Stub1* were equal across all three genotypes (Figure 3B). We isolated protein extracts from patients homozygous for the CHIP-T246M mutation, and again, we measured a decrease in steady-state soluble CHIP-T246M protein (Figure 3C). Together these data suggest that the reduction in CHIP-T246M protein is likely a result of post-translational regulation.

**Figure 3:**
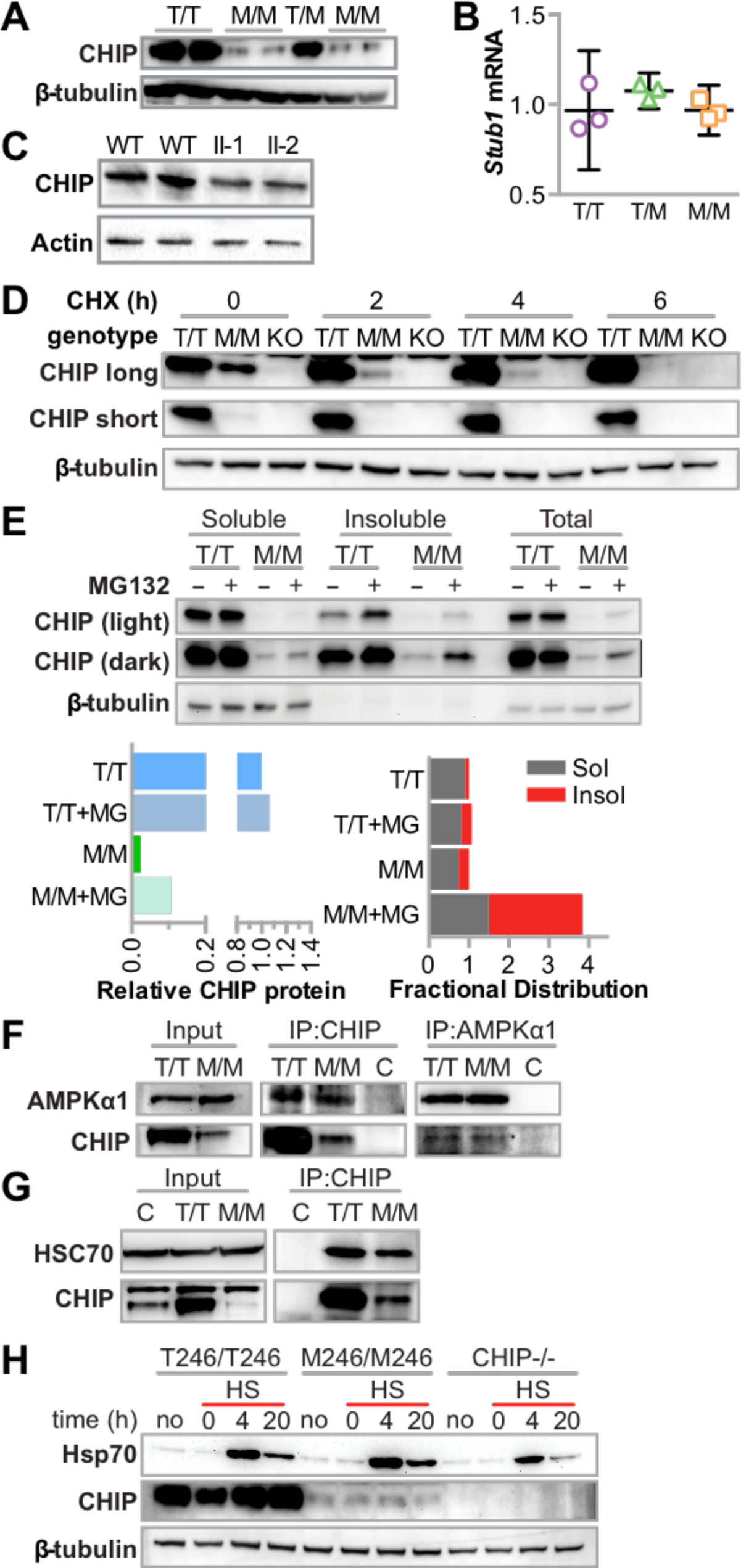
CHIP-T246M expressed from the endogenous locus results is regulated post-translationally resulting in decreased steady-state protein levels, however, interactions between CHIP and known interactors are not affected. (**A**) Immunoblot analysis of CHIP and β-tubulin protein or (**B**) qPCR analysis of *Stub1* mRNA levels in primary fibroblasts isolated from T246/T246 (T/T), T246/M246 (T/M), or M246/M246 (M/M) mouse embryos. Relative mRNA levels are represented by the dot plot and summarized by the mean and 95% confidence intervals. (**C**) Immunoblot analysis of CHIP and β-actin protein in fibroblasts isolated from control patients (WT) or siblings that are homozygous for CHIP-T246M (II-1 and II-2). (**D**) Immunoblot analysis of CHIP and β-tubulin in soluble cell lysates from either T/T or M/M, fibroblasts, or fibroblasts isolated from CHIP-/-embryos (KO) after exposure to cycloheximide (CHX) indicated in hours (h). Two exposures of CHIP immunoblots are provided to help visualize M/M conditions. (**E**) Immunoblot analysis of CHIP and β-actin in either the soluble or insoluble fraction of lysates or whole cell lysates from T/T or M/M MEFs treated with 20 µM MG132 or 0.05% DMSO control for 4 hours. (**F**) Immunoblot analysis of AMPKα1 and CHIP in cell lysates from T/T or M/M MEFs either before (input) or after immunoprecipitation (IP) of the indicated protein. Control samples (C) contained a mixture of 50% T/T and T/M and were immunoprecipitated with either rabbit IgG or goat IgG as controls for the CHIP and AMPKα1 antibodies, respectively. (**G**) Immunoblot analysis of HSC70 and CHIP in cell lysates from T/T or M/M MEFs either before (input) or after immunoprecipitation (IP) of CHIP. Control samples (C) contained a mixture of 50% T/T and T/M and were immunoprecipitated with rabbit IgG to control for the CHIP antibody. (**H**) Immunoblot analysis of HSP70 and CHIP in MEFs with the indicated genotypes that were treated without heat shock (no) or with heat shock (HS) followed by the indicated recovery time.

#### Endogenous CHIP-T246M is degraded by the proteasome

Given the difference in steady-state soluble protein levels of CHIP-T246M in primary cells from patients and our preclinical models, we evaluated the turnover rate of CHIP in MEFs. Similar to exogenously expressed CHIP proteins (Figure 2G), the turnover of soluble CHIP-T246M protein was rapid (t1/2 = 1.2 h) compared to the stable, CHIP-WT protein. In fact, CHIP-T246M protein was undetectable after 6 hours of cycloheximide chase compared to approximately 75% of CHIP-WT protein that remained after 6 hours (Figure 3D). To evaluate the solubility distribution of CHIP-T246M and whether this distribution was effected by proteasome inhibition, we treated MEFs with 20 µM MG132 or 0.05% DMSO control for 4 hours. We collected soluble, insoluble and total protein fractions from samples containing equal cell numbers and analyzed protein levels using immunoblot analysis. Total T246M protein is dramatically reduced compared to WT protein (Figure 3E). Secondly, the change in total protein levels is dramatically higher with proteasome inhibition for T246M (4-fold) protein compared to WT, suggesting a robust proteasome-dependent turnover of endogenous CHIP-T246M (Figure 3E).

#### CHIP-T246M maintains protein-protein interactions and the response to heat shock

We previously demonstrated that exogenous CHIP-T246M immunoprecipitates more chaperone clients, such as HSP70 and HSC70, compared to CHIP-WT [11]. However, given the strong decrease in steady-state CHIP-T246M protein expression seen in primary cells, it is possible that these interactions are lost, resulting in a CHIP null phenotype. Remarkably, in primary cells we found that CHIP-T246M maintains interactions with its chaperone substrate AMPK (Figure 3F), as well as HSC70 (Figure 3G), at levels similar to CHIP-WT, suggesting that CHIP-T246M may still exhibit activity towards these proteins. To test this concept, we exposed primary cells to heat and to determine if cells expressing CHIP-T246M still respond to heat shock, as CHIP plays an important role in inducing the heat shock response via activation of HSP70. Unexpectedly, M246/M246 cells still maintained activation of HSP70 during the recovery period following heat shock (Figure 3H), whereas cells completely lacking CHIP expression had an attenuated response as previously described [27]. These data demonstrate that CHIP-T246M may retain some chaperone function, despite the increase in turnover.

### PRECLINICAL MODELS OF CEREBELLAR CHIPOPATHY

#### Generating in vivo models of CHIP-T246M

Given the advantages of both mouse and rat models to study neurological disease, we developed a rat model also harboring the same endogenous CHIP-T246M mutation. Similar to the decrease seen in fibroblasts isolated from patients with T246M mutations, steady-state levels of CHIP-T246M expression are 40% lower in protein extracts isolated from the cerebellums, whole brain, and testes of T246M rats (Figure 4A) and further decrease with age (Figure 4B). CHIP protein levels in mouse tissues isolated from M246 mice were more dramatically decreased (90% decreased) compared to CHIP levels in tissues from wild-type mice (Figure 4C). The decrease in tissue expression was not due to differences in levels of the mRNA that encodes CHIP (Figure 4D). The effect of the T246M mutation on CHIP was also evident via immunohistochemical analysis in both the mouse and rat models (Figure 4E) where we also identified Purkinje cell degeneration, evidenced by a loss in calbindin staining (Figure 4E). In both mouse and rat models, we observed lower body weights in the M246/M246 mice over time (Figure 4F) as well as an increase in mortality in rats (Figure 4G). In addition to lower body weights, there was selective tissue atrophy in the brain and testes of M246/M246 mice, known tissues that play a role in SCAR16 pathology, whereas the heart was not affected (Figure 4H). Together, these data demonstrate that our two preclinical models of T246M recapitulate features of cerebellar CHIPopathy in patients including decreased protein expression of CHIP, neurodegeneration, as well as neuro- and gonadal-atrophy.

**Figure 4:**
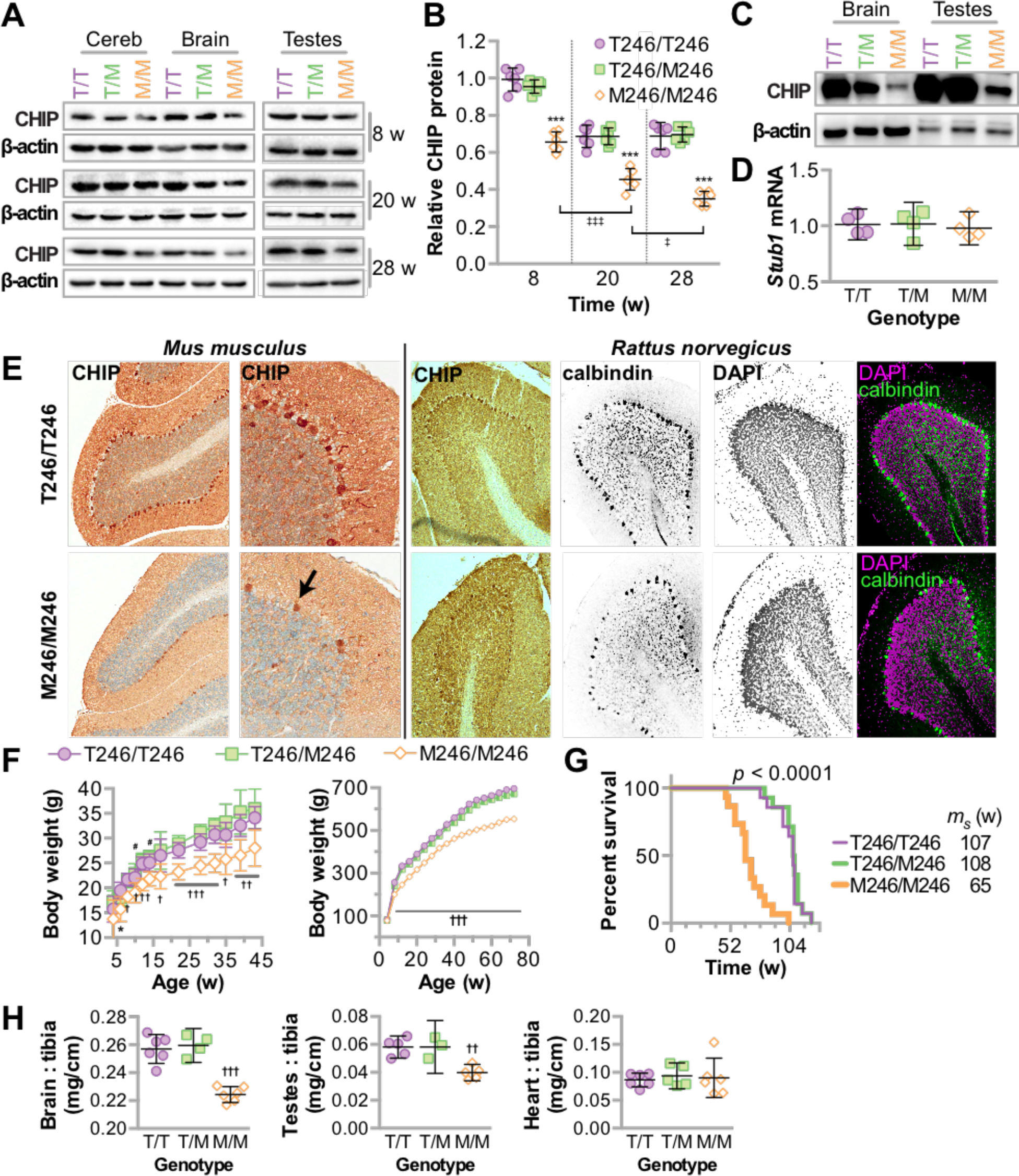
Rodents engineered with CHIP-T246M had decreased expression of CHIP in the brain and testes, Purkinje cell degeneration, increased mortality, and selective tissue atrophy. (**A**) Immunoblot analysis of CHIP and β-actin in the cerebellum (cereb), brain, and testes of rats with either T246/T246 (T/T), T246/M246 (T/M), or M246/M246 (M/M) genotypes at harvested at different ages. (**B**) Densitometry analysis of relative CHIP protein represented by dot plot and summarized by the mean ± 95% CI: *** *p* < 0.001 compared to T/T or T/M; ‡, ‡‡‡ correspond to *p* < 0.05, or 0.001 compared to previous time point via Tukey’s post hoc test. (**C**) Immunoblot analysis of CHIP and β-actin in the brain and testes of mice with the indicated genotypes. (**D**) qPCR analysis of *Stub1* mRNA levels in four different tissues (heart, liver, brain, testes) isolated from mice with the indicated genotypes, n = 3 mice per tissue. Relative mRNA levels are represented by the dot plot and summarized by the mean and 95% confidence intervals. (**E**) Representative micrographs of immunohistochemical detection of CHIP expression or indirect immunofluorescence of calbindin expression in cerebellums of mice (*Mus musculus*) and rats (*Rattus norvegicus*) with the indicated genotypes. DAPI is included for a fluorescent counterstain. The arrow indicates a cell that retains CHIP expression. (**F**) Total body weight of mice (left) or rats (right) with the indicated genotypes over age, Tukey’s post hoc test: * p < 0.05 M/M vs. T/T; †,††, ††† correspond to *p* < 0.05, 0.01, 0.001 comparing M/M to T/T and T/M; # *p* < 0.05 M/M vs. T/M. (**G**) Survival analysis of rats with the indicated genotypes (Mantel-Cox test) with the median survival (m_s_) indicated in weeks (w). Tissue weights normalized to tibia length in mice with indicated genotypes represented by dot plot and summarized by the mean ± 95% CI: ††, ††† correspond to *p* < 0.01, 0.001 comparing M/M to T/T and T/M.

#### M246 results in progressive ataxia and alterations in gait

One measure of ataxia in rodent models is measured by performance on a rotating rod, known as the rotarod test. In both mouse and rat models, we found a decrease in rotarod performance. In mice, we found initial learning to be blunted (Figure 5A) and an age-dependent decrease in performance starting around 30 weeks of age (Figure 5B). Given our previous observation of a near complete lack of learning of rotarod behavior in mice completely lacking CHIP expression [11], we utilized a second metric of ataxia progression. This composite test is comprised of hind limb clasping, ledge test, gait, and kyphosis [29] and was used to measured ataxia onset and progression in both the CHIP-T246M and CHIP(-/-) mouse lines (Figure 5C). Remarkably, we found that CHIP-/- mice were already ataxic at weaning (score = 4.4, **Video 1**), and the rate of ataxia progression was equivalent to wild-type mice (rate = 0.17 vs. 0.20 points/week in -/- vs. +/+ mice, respectively). In contrast, M246/M246 mice had a high rate of disease progression, 0.38 point/week (**Video 2A**), suggesting the T246M mouse may reflect a more suitable preclinical model for SCAR16 compared to the CHIP-/- mouse. Likewise, in rats, we found robust decreases in rotarod performance at ages as young as 12 weeks of age that rapidly decreased with age (Figure 5D and **Video 2B**). The loss in rotatrod performance was also accompanied by changes in gait (Figure 5E-5H), and much like the effect on motor performance, the effect on gait was exacerbated by age with notable differences occurring at 32 weeks of age. Thus both the mouse and the rat model of CHIP-T246M demonstrate progressive ataxia that worsens with age and recapitulates the clinical phenotype observed in patients with cerebellar CHIPopathy that suffer from an early adult-onset progressive ataxia [12].

**Figure 5:**
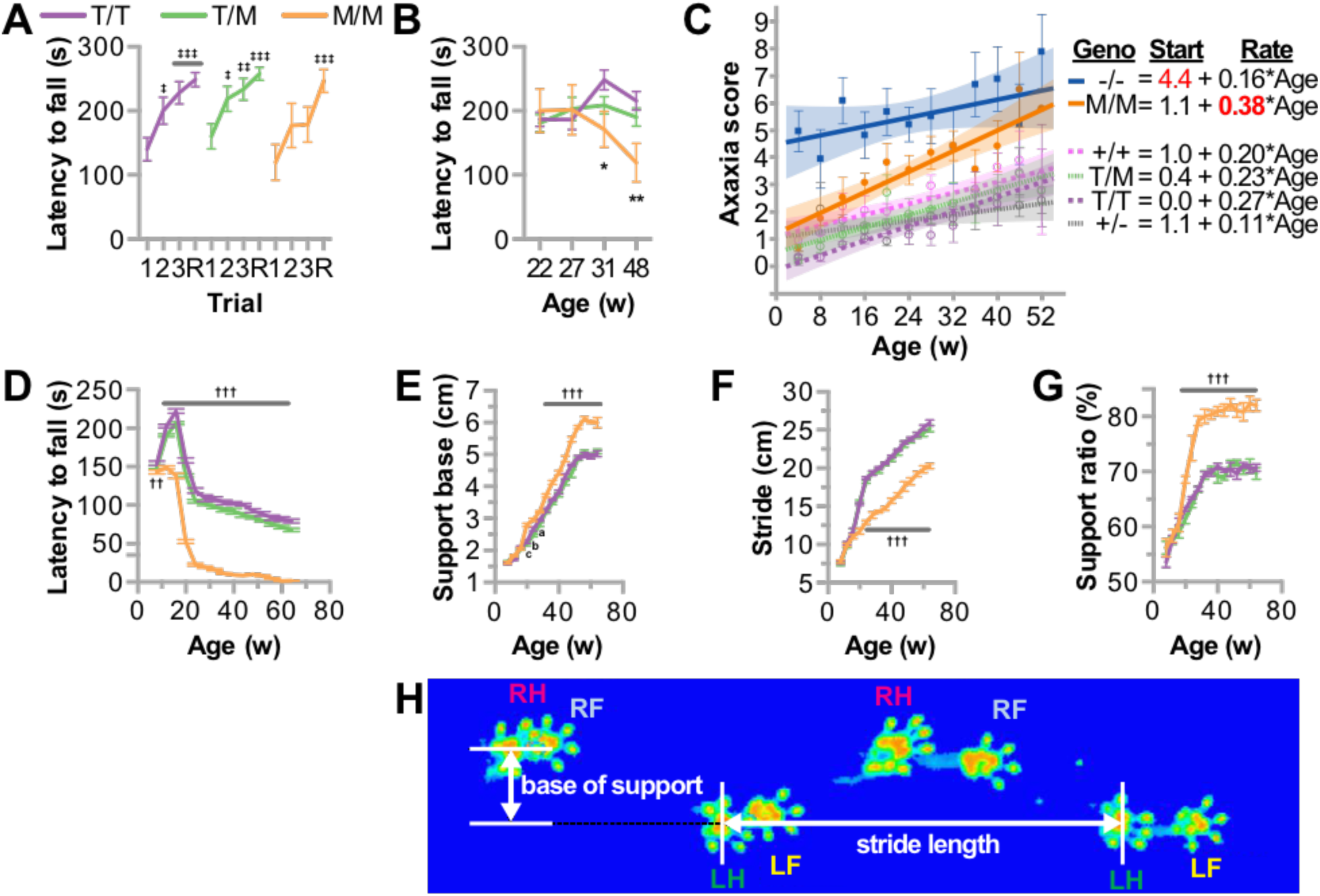
CHIP-T246M in rodents resulted in progressive ataxia and altered gait. Rotarod analysis during the initial learning phase (**A**) or over time (**B**) represented by line graph and summarized by the mean ± SEM (N = 9-20 animals/genotype), Tukey’s post hoc test: ‡, ‡‡, ‡‡‡ correspond to *p* < 0.05, 0.01, 0.001 compared to previous time point; *, ** correspond to *p* < 0.05, 0.01 comparing M/M vs. T/T. (**C**) Composite ataxia score of mice with the indicated genotypes including wild-type (+/+), heterozygous (+/-), and homozygous (-/-) knockouts of the CHIP allele represented by scatterplot and summarized by the mean ± SEM (N = 12 animals/genotype). Results of the linear regression analysis are depicted by the line and equation provided. Changes in (**D**) rotatrod performance, (**E**) support base, (**F**) stride, and (**G**) support ratio over time in rats with the indicated genotypes represented by line plot and summarized by the mean ± SEM (N = 6 animals/genotype), Tukey’s post hoc test: a, b, c, and ††† correspond to *p* < 0.05, 0.01, 0.001, and 0.001 comparing M/M to T/T and T/M. (**H**) Graphical representation of gait parameters with feet locations indicated (RH = right hand, RF = right foot, LH = left hand, LF = left foot).

#### Behavioral repercussions of CHIP-T246M include loss of prepulse inhibition, hyperactivity, and cognitive dysfunction

We utilized our rodent models to determine effects of the T246M mutation on the additional clinical hallmarks of SCAR16, decreased sensorimotor reflexes and cognitive dysfunction. Given the onset of ataxia phenotype in both the mouse and rat models were observed around 30 weeks of age (Figure 5B, 5C), we tested mice at ages both before and after this time point, 8-12 and 33 weeks, respectively. Using the acoustic startle and prepulse inhibition test we found no effects of M246 on the amplitude of the acoustic startle response in either young or adult mice (Figure 6A). Similarly, all young mice had comparable levels of prepulse inhibition; however, older M246/M246 mice exhibited robust decreases in percent inhibition, indicating the emergence of sensorimotor deficits by age 33 weeks (Figure 6B). These degenerative deficits in prepulse inhibition are consistent with other mouse models of cerebellar degeneration with profound Purkinje cell loss [30].

**Figure 6:**
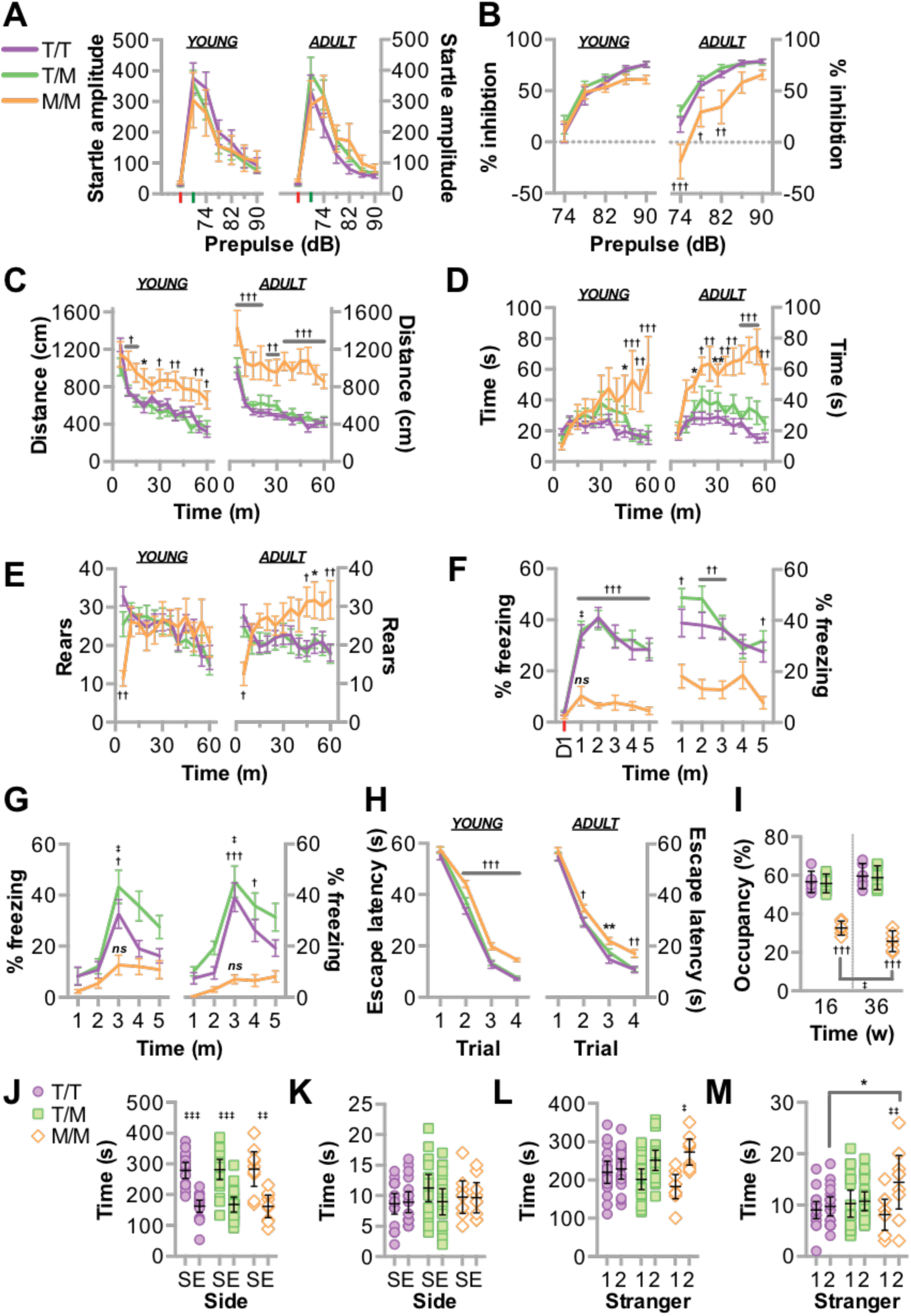
CHIP-T246M resulted in age-dependent deficits in sensorimotor skills and increased anxiety; rodents with CHIP-T246M also developed cognitive deficits and changes in sociability. (**A**) The amplitude of startle response (red and green are no stimulus and acoustic startle stimulus alone trials, respectively) and (**B**) the amount of prepulse inhibition over increasing levels of sound, decibels (dB). (**C**) The distance traveled, (**D**) time spent in the center, and (**E**) the number of rearing movements in open field test over time. Tests (**A-E**) were performed on mice with the indicated genotypes at 11-13 w of age (young) and repeated at 33 w of age (adult). Conditioned fear testing is calculated by the percent freezing during either (**F**) context or (**G**) cue-dependent learning during the training day (D1), and over 5 minute periods either a day after training (left) or two weeks later (right). Tests **A-G** are represented by line plot and summarized by the mean ± SEM (N = 10 – 20 animals/genotype), Tukey’s post hoc test: *, ** correspond to *p* < 0.05, 0.01 comparing M/M vs. T/T; †,††, ††† correspond to *p* < 0.05, 0.01, 0.001 comparing M/M to T/T and T/M; ‡ corresponds to *p* < 0.001 compared to the initial stimulus within the genotype (learning), ns = *p* > 0.05. (**H**) The average latency to find the platform over four consecutive training days in the Morris Water maze task is represented by line plot and summarized by the mean ± SEM. (**I**) The amount of time spent in the target quadrant when the platform was removed after the last trial is represented by dot plot and summarized by the mean ± 95% CI. Tests **H-I** were administered on rats with the indicated genotypes at 16 w of age (young) and repeated at 36 w of age (adult), N = 6 animals/genotype, Tukey’s post hoc test: ** corresponds to *p* < 0.01 comparing M/M vs. T/T; †,††, ††† correspond to *p* < 0.05, 0.01, 0.001 comparing M/M to T/T and T/M; ‡ corresponds to *p* < 0.05 within genotype. (**J**) Time spent and (**K**) number entries in either the side with stranger 1 (S) compared to an empty cage (E). (**L**) Time spent and (**K**) number entries in either the side with previous stranger 1 (1) or novel stranger 2 (2). Tests **J-M** were performed on mice with the indicated genotypes represented by dot plot and summarized by the mean ± 95% CI (N = 10-20 animals/genotype), Tukey’s post hoc test: ‡, ‡‡, ‡‡‡ correspond to *p* < 0.05, 0.01, 0.001 within genotype; * p < 0.05 M/M vs. T/T.

Mice were tested in the elevated plus maze at eight weeks of age and there was no difference between genotypes in any of the parameters measured (Table 2). However, we observed age-dependent defects including increases in hyperactivity and impulsivity in an open field test. The M246/M246 mice traveled father (Figure 6C) and spent more time in the center of the field (Figure 6D) compared to control mice or when the mice were younger. Interestingly, a different pattern emerged for rearing movements, a measure of vertical activity, as the M246/M246 mice had reduced levels of rearing at the beginning of each session (Figure 6E), indicating a deficit in the initial exploration of the open field, however, in older mice, higher levels of rearing emerged in the mutant group in the last half of the session, consistent with a hyperactive phenotype (Figure 6E).

**Table 2:** Elevated plus maze, marble burying, and olfactory sensing in CHIP-T246M mice. The performance measurements from the elevated plus maze test, the marble-burying assay, and buried food test for olfactory function are represented by the mean ± SEM from mice with the indicated genotypes, † p < 0.05 comparing M/M to T/T and T/M.

|  | T246/T246 | T246/M246 | M246/M246 |
| --- | --- | --- | --- |
| **Elevated plus maze** |  |  |  |
| Percent open arm time | 32 ± 3 | 30 ± 4 | 36 ± 4 |
| Percent open arm entries | 39 ± 2 | 41 ± 2 | 42 ± 3 |
| Total number of entries | 28 ± 2 | 27 ± 2 | 28 ± 3 |
| **Number of marbles buried** |  |  |  |
| First test (age 12 w) | 16 ± 0.6 | 18 ± 0.4 | 11 ± 1.7 $^{\dagger}$ |
| Second test (age 32 w) | 17 ± 0.4 | 16 ± 0.7 | 13 ± 1.3 $^{\dagger}$ |
| **Olfactory test** |  |  |  |
| Latency to find food (s) | 267 ± 74 | 189 ± 52 | 289 ± 95 |
| Percent of group finding food | 80% | 95% | 89% |

Our data suggest CHIP-T246M results in impulsive and risky exploration, as observed in mouse models for mania and impulsivity and overt hyperactivity [31, 32]. Interestingly, both impulsivity and hyperactivity have been attributed to cognitive cerebellar dysfunction in humans [33–35]; therefore, we evaluated cognitive function in mice using the fear response test and in rats using the Morris water maze. The M246/M246 mice had impairments in the conditioned fear procedure, both in contextual and cue-dependent learning (Figure 6F, 6G).

On the initial training day, all genotypes had similar, low levels of freezing before any exposure to the aversive foot shock. On the next day, or two weeks later, the lack of learning in M246/M246 mice was apparent as they did not increase their freezing time to either a contextual (Figure 6F) or cue-dependent (Figure 6G) stimulus. This lack of response could not be attributed to hearing impairment, since the mutant mice had normal performance in the acoustic startle test (Figure 6A). Finally, to determine if the M246 mutation has similar effects on cognition in the rat model, we measured the performance of young and adult rats in the Morris water maze. Consistent with the conditioned fear responses in mice, learning was also impaired in M246/M246 animals, as measured by the escape latency (Figure 6I). Moreover, young M246/M246 rats had a 42% decrease platform zone occupancy that worsened in older animals (Figure 6H), consistent with a profound deficit in memory recall. Together these data suggest significant impairment in learning and memory as a result of CHIP-T246M and are consistent with the clinical phenotype of patients with T246M [11].

Additional testing in the 3-chamber choice test found that the M246/M246 genotype associated with altered social behavior (Figure 6J**-6M**), such that M246/M246 mice had increased preference for social novelty towards the newly-introduced stranger 2 (Figure 6L, 6M). Further behavioral testing revealed the M246/M246 mice had reduced marble burying, indicating a decrease in exploratory digging (Table 2); moreover, no effects of genotype were observed for olfactory ability in a buried food test (Table 2). Overall, the results of the battery of behavioral assessments performed suggest that homozygous CHIP-T246M leads to the dysregulation of inhibitory processes governing activity, exploration, and sensorimotor gating, as well as impaired learning and memory in tests for conditioned fear and cognition. Interestingly, impaired conditioned fear and decreased marble-burying were reported in mice with deletion of maternal E3 ubiquitin ligase *Ube3a*, a model for Angelman syndrome [36], again highlighting the critical role of protein ubiquitination in cerebellar homeostasis.

#### Alterations in the T246M proteome identified both known and potentially novel substrates of CHIP-dependent regulation

Given the role of CHIP as a chaperone, co-chaperone, and ubiquitin ligase it is likely that disruption in CHIP function alters the regulation of proteins critical to cerebellar function and protein homeostasis. To identify candidate proteins that may mediate the pathogenesis of SCAR16, we performed unbiased proteomics via mass spectroscopy to identify differentially expressed proteins in cerebellar lysates prepared from either T246/T246 or M246/M246 rats. We identified 63 and 80 unique proteins that were either less or more abundant in M246/M246 relative to T246/T246 cerebellums (Figure 7A, **Supplementary Figure S1, Supplementary Table S1**). As expected, we identified a 50% decrease in CHIP protein in M246/M246 brains as measured via MS (mean ratio = 0.5, *p* = 3.47E-08). We confirmed the differential levels of selected proteins that were either increased in M246/M246 cerebellums including Pde9a and Tau or decreased in M246/M246 cerebellums, such as Phlpp1, alpha-synuclein, CHIP, and Pink1 (Figure 7A). Next, we analyzed the dataset using known and predicted protein-protein association data combined with functional enrichment analysis via the STRING database [37] to identify proteins or pathways that may play a role in the cerebral pathophysiology of SCAR16. Of the 143 initial proteins, 60 proteins were identified in a total of 21 protein-protein interaction clusters (**Supplementary Table S2**) including a family of six proteins that contained CHIP and other regulators of protein quality control, such as BAG3 and several F-box proteins (Figure 7B). Additionally, we identified a cluster of proteins known to be affected by CHIP expression including tau (MAPT), alpha-synuclein (SNCA), and PINK1 [5, 38, 39]. Of note, seven member cluster that was also functionally enriched for coagulation processes (wound healing) and extracellular matrix function was identified (Figure 7B), and in total, 16 proteins involved in coagulation were affected by CHIP-T246M. Coagulation was recently identified as a significant component of spinocerebellar ataxia disease signatures (without Friedreich ataxia) from a large-scale analysis of 80,000 samples [40] and our analysis of CHIP-T246M cerebellums support the notion that coagulation may represent new targets for therapeutic interventions.

**Figure 7:**
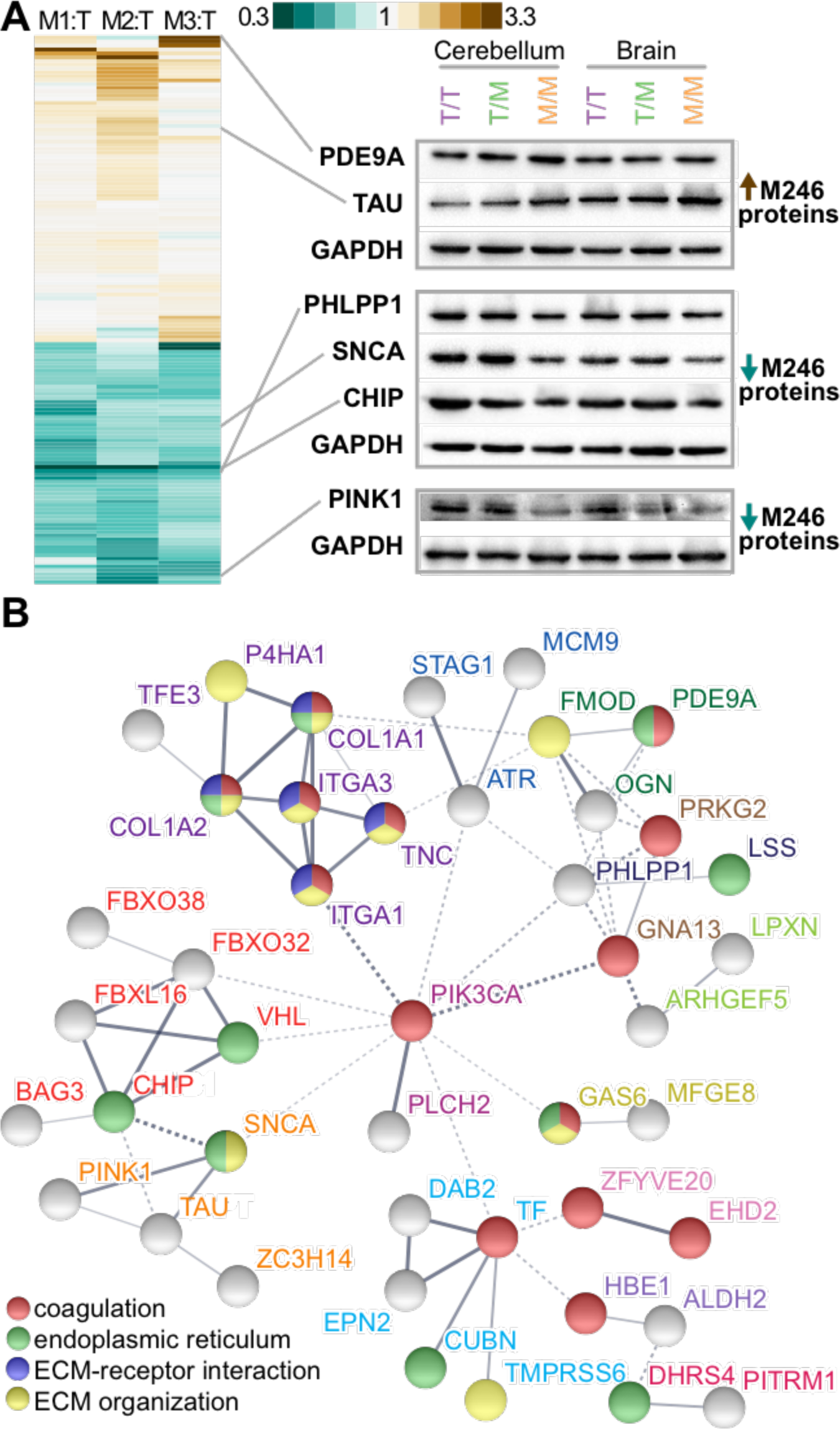
Changes in the cerebellar proteome due to CHIP-T246M. (**A**) Heatmap (**left**) of cerebellar proteins determined to be differentially expressed in M246/M246 relative to T246/T246 rats and immunoblot analysis (**right**) of selected proteins in either cerebellums or whole brain in rats with the indicated genotype, GAPDH was used as a loading control. (**B**) Protein interactions analysis identified both the interaction clusters and functional enrichment of the differentially expressed proteins identified due to CHIP-T246M. The interaction clusters are connected with solid lines and additionally indicated by colored protein labels. Functional enrichment analysis is labeled by the colored spheres.

## Discussion

CHIP plays a vital role in cerebellar homeostasis, however, how do the various mutations that cause cerebellar CHIPopathy drive the disease pathogenesis? The simplest explanation is that the CHIP mutations destabilize protein stability that lead to an effective CHIP null condition. However, a varying degree of CHIP protein levels is found in fibroblasts isolated from patients with CHIP mutations. For example, the N65 mutation results in an 80% loss of CHIP protein [21], whereas other mutations, such as T246M (Figure 3C) and the compound mutation M211I/E238* have more modest effects, approximately at the level of haploinsufficiency [16]. Moreover, the R119*/I294F and K145Q/P243K compound mutations do not appear to affect CHIP turnover [20]. Given the recessive nature of this disease in patients [12], and the lack of phenotype observed in our three rodent models of CHIP haploinsufficiency (Figure 5C, 5D), these data suggest that either the loss or change in specific activities of CHIP drive the pathophysiologies associated with cerebellar CHIPopathy.

We identified that the T246M leads to disruption of the U-box structure (Figure 1A, 1B, 1C), and the effect of the T246M conformational change on CHIP function appears to be three-fold. First, the destabilization of the U-box results in no appreciable ligase activity as demonstrated previously by our group [11] and others [23, 24]. Secondly, CHIP-T246M is degraded in a proteasome-dependent manner at a higher rate than CHIP-WT, both when expressed exogenously (Figure 2G) and under native genomic conditions (Figure 3D, 3E), resulting in decreased steady-state levels that we observed both in both T246M patients (Figure 3C) and in our rodent models (Figure 4A, 4B, 4C). Lastly, and more surprisingly, the T246M mutation promotes the formation of soluble oligomeric forms of CHIP comprised of 10-12 mers. CHIP functions as a dimer [41, 42], however under heat stress, CHIP forms higher-order oligomers that appear to have increased chaperone activity [8]. Therefore, we tested the effect of T246M on the chaperone-mediated activation of HSF1 and found that T246M results in increased nuclear localization of HSF1 (Figure 2A) and increased HSF1 activity (Figure 2B). We initially considered that CHIP-T246M was activating HSF1 simply because the cells were responding to a misfolded protein insult, and not a specific effect of CHIP-T246M on HSF1. To account for this, we engineered a CHIP-T246M protein that also contained the K30A mutation that abolishes the interaction between CHIP and its chaperone binding partners. Unlike T246M, the K30a-T246M double mutant had little effect on HSF1 activity (Figure 2B) despite similar levels of insoluble CHIP protein (Figure 2F). Therefore, while the T246M may abolish ligase activity, it appears that one consequence of these mutations may be altering chaperone function by failing to process substrates usually triaged by the CHIP-HSC(P)70 complex in a manner distinct from the CHIP null condition. We previously observed increased pulldown of HSP70 and HSC70 with CHIP-T246M [11], and even in our rodent model with reduced steady-state levels of CHIP-T246M, this interaction is maintained (Figure 3G). In fact, comparable amounts of HSC70 as well as the chaperone substrate of CHIP, AMPK, immunoprecipitate with CHIP or CHIP-T246M (Figure 3F, 3G) suggesting that even though the expression of CHIP-T246M is reduced, it is still engaging with known client proteins at similar levels. In cell culture models, the synthetic CHIP mutation, H260Q, acted in similar to T246M regarding HSF1 activation (Figure 2B), confirming a previous report that also identified that a U-box mutation led to changes in BAG3 protein levels and suppression of the macro autophagy pathway [43]. Likewise, it was proposed that enhancing either macro or chaperone mediated autophagy may be beneficial in CHIPopathies [16]. Interestingly, our proteomics analysis of the T246M rodent cerebellum revealed a decrease in Bag3 protein levels (**Supplementary Figure S1, Supplementary Table S1**), so one distinct possibility for the pathology associated with cerebellar CHIPopathy may be a disruption in either the macroautophagy or chaperone-mediated autophagy pathways, rendering cells susceptible to proteinopathies. Alteration of these autophagy pathways is implicated in other ataxias, such as spinocerebellar ataxia type 1, 3 7, and 14, [44–47].

Central to disease-based research is the establishment of preclinical models that can be used as a platform to both better understand the mechanism of disease and to test therapies. Therefore, we created both a mouse and rat model that genocopies the human T246M. We found that these models recapitulate the key features of SCAR16, notably: decreased steady-state protein expression due to proteasome-dependent turnover (Figure 3D, 3E, 4A, 4B, 4C, 4E); tissue atrophy (Figure 4H); increased mortality (Figure 4G); loss of motor function (Figure 5C, 5D); age-dependent impairment of sensorimotor function (Figure 6B), overt hyperactivity (Figure 5C, 5D, and 5E); and impaired cognitive function (Figure 6F, 6G, 6H, 6I). Given these phenotypes were exacerbated with age, these models provide us with the ability to interrogate the pathophysiology at difference time points in disease progression. These data further support our previous findings that CHIP plays a critical role in cerebellar maintenance [11]. Interestingly, while some behavioral deficits in M246/M246 rodents overlap with those observed in CHIP-/- mice [11], the majority were unique to T246M mutation, further supporting our hypothesis that disease-causing mutations in CHIP and the total loss of CHIP are not functionally equivalent. For example, M246/M246 mice exhibited overt hyperactivity (Figure 6C) risky behavior (Figure 6D), deficits in prepulse inhibition (Figure 6B), and increased preference for social novelty (Figure 6K, 6M), phenotypes not observed in CHIP-/- mice [11]. We hypothesize that the phenotypic differences observed between CHIP-/- mice and T246M mice are likely reflective of our cell-based and *in vitro* findings that while T246M CHIP no longer functions as an E3 ubiquitin ligase, other CHIP functions remain intact despite this mutation.

Perhaps even more intriguingly, the T246M mutation may modify the co-chaperone activity of CHIP in a deleterious manner. For example, mutant CHIP may be unable to either ubiquitinate chaperone-engaged proteins or promote the refolding of proteins that are usually degraded by the proteasome. Similarly, mutant CHIP may promote the activation of compensatory pathways involved in proteins degradation or clearing protein aggregates such as autophagy [43] that could sensitize the cells to additional proteotoxic stress. Moreover, the formation of soluble CHIP oligomers that alter the chaperone or co-chaperone functions of CHIP may also contribute to the pathophysiology. Remarkably, the oligomerization of CHIP-WT with heat is thought to increase its intrinsic chaperone activity by enhancing the binding activity of CHIP to substrates [8], thereby providing an additional mechanism through which coding mutations could affect CHIP activity. Alternatively, CHIP could function as a sink for other components of the ubiquitin proteasome system, such as E2 enzymes, ultimately altering the fate of proteins that are usually modified either by CHIP or other E3 enzymes that share the same E2 enzymes. In total, our data support our previous findings that highlight the role of aberrant ubiquitination in the pathogenesis of SCAR16; however, the loss of CHIP ubiquitin ligase activity alone may not fully explain the molecular mechanisms underlying the diverse pathophysiology observed in the heterogeneity of SCAR16 disease.

CHIP mutations associated with SCAR16 occur in all three of CHIP’s functional domains, although interestingly the majority are concentrated in the charged domain and the U-box domain. Biochemical studies using in vitro approaches demonstrated wide variance regarding how individual properties of CHIP change given the location of the mutation [23, 24]; clearly, more cell-based and in vivo approaches are needed to fully understand how these mutations drive a disease that has a spectrum of clinical phenotypes. Given the diversity of mutations and the clinical heterogeneity of the ARCA patients harboring *STUB1* mutations, we originally hypothesized that the affected protein domain might directly correlate to clinical phenotype. For example, cognitive impairment occurred in all SCAR16 patients described in the literature that harbor mutations in the U-box domain [11, 15, 17–20, 22], such that residual CHIP activity involving a defective or truncated U-box domain but intact TPR domain could directly correlate to specific clinical symptoms in some patients. Our extensive behavioral analysis of two T246M mammalian models provides direct evidence to support this hypothesis, demonstrating that particular cognitive deficits are in fact associated *in vivo* with a U-box domain point mutation that has been demonstrated *in vitro* and in cells to have a partially functionally intact TPR domain. The development of additional animal models with specific domain mutations may help to further validate this hypothesis and identify how the multifunctional roles of CHIP contribute to particular clinical pathologies. It seems clear, however, that while defective ubiquitination contributes to SCAR16 pathology, CHIP mutation as a driver of disease is not limited to loss of ubiquitin ligase activity but may represent a more multi-faceted disruption of CHIP-mediated PQC.

In light of the identification of T246M CHIP mutation and subsequent designation of SCAR16, the establishment of preclinical models of SCAR16 represents novel and important tools to evaluate CHIP dysfunction *in vivo* in a disease-relevant context. Our biophysical, cellular and *in vivo* characterization of T246M mutation provides significant insight into both the molecular mechanisms driving disease pathology in SCAR16 as well as basic CHIP biology, by shedding new light on the structure-function relationship, particularly regarding the multifaceted activities of CHIP within the cell. Furthermore, we are hopeful these studies provide valuable insight required for the future development of effective therapies for this devastating degenerative disease.

## Experimental Procedures

### BIOCHEMICAL AND CELLULAR STUDIES

#### Expression plasmids and recombinant proteins

Mammalian expression plasmids pcDNA3-myc-CHIP, pcDNA3-myc-CHIP-K30A, pcDNA3-myc-CHIP-H260Q, HA-Ubiquitin, FLAG-SIRT6, FLAG-HSP70, β-galactosidase, and GFP were used as described previously [3, 8, 43, 48, 49]. CHIP, CHIP-H260Q, CHIP-K30A, CHIP-T246M and AMPK recombinant proteins were produced in Escherichia coli BL21(DE3) as His-tagged fusion proteins by induction with 0.1mM isopropyl-1-thio-β-D-galactopyranoside overnight at 18°C followed by purification with HisTrap™ HP columns (GE Healthcare), concentrated, and stored in in 20 mM HEPES pH 7.4 with 150 mM NaCl.

#### Mutagenesis

A point mutation of threonine to methionine at position 246 of CHIP was created for generation of single T246M point mutant and K30A-T246M double point mutant by site-directed mutagenesis using the Q5 Site-Directed Mutagenesis Kit (New England Biolabs, E0554S) according to manufacturer’s instructions using previously described pcDNA3-myc-CHIP template or pcDNA3-myc-K30A CHIP template [3] and mutagenic primers 5’-CCGTGCATCATGCCCAGTGGC-3’ and 5’-CTCCCGCATCAGCTCAAAGC-3’ (BaseChanger software, New England Biolabs). The myc-CHIP-T246M and myc-CHIP-K30A-T246M expression plasmids were produced by transformation in Escherichia coli DH5α, purified, and the single-base pair substitution was verified by DNA sequencing.

#### Light scattering

The solution molecular weights of WT CHIP, K30A, H260Q and T246M point mutant CHIP were determined by size exclusion chromatography followed by multi-angle light scattering (SEC-MALS) of the eluant from a size exclusion chromatography column. The SEC-MALS system consisted of a GE Superdex 200 column connected to Wyatt DAWN HELEOS-II multi-angle light scattering instrument and a Wyatt T-Rex refractometer (Wyatt Technology, Santa Barbara, CA, USA). 100 µl of 0.5 mg/ml of each sample was loaded onto the column, and the light scattering and refractive index data were collected as the eluted samples passed through light scattering system. The molar masses of the samples eluting in various peaks were calculated from these data using ASTRA 6 software (Wyatt Technology).

#### Nuclear magnetic resonance spectroscopy

Human WT and T246M U-box (amino acid residues 212-303) recombinant proteins were expressed and purified as previously described for WT CHIP U-box [50]. NMR spectra were recorded using a Bruker Avance 600 MHz (1H) spectrometer at 20 °C in buffer containing 20 mM HEPES (pH 7.5), 50 mM NaCl, and 1 mM DTT as previously described [50]. NMR data were processed with NMRPipe [51] and analyzed with SPARKY (T. D. Goddard and D. G. Kneller, SPARKY 3, University of California, San Francisco).

#### Circular dichroism spectroscopy

CD spectra of WT and T246M U-box CHIP were collected as previously described [52] at 0.25 mg/mL and 15 °C in 10 mM sodium phosphate (pH 7.0) with 20 mM NaCl and 1 mM DTT. The Tm for WT and T246M U-box CHIP was determined by monitoring the temperature dependence of CD at 222 nm.

#### Cell culture and transfection

CHIP^+/+^, CHIP^−/−^ and T246M CHIP mouse embryonic fibroblasts (MEFs) were cultured as previously described [27]. COS-7 and shCTRL and shCHIP HEK293 cells were maintained in Dulbecco’s modified Eagle’s medium (Invitrogen) supplemented with 10% fetal bovine serum (Sigma) at 37 °C in an atmosphere of 5% CO_2_. Cell transfections were performed using X-tremeGENE 9 (Roche) with the indicated plasmid DNA at a 1:3 DNA to X-tremeGENE 9 ratio.

#### mRNA analysis

CHIP mRNA levels in primary MEFs or tissue were determined using the SingleShot™ SYBR® Green Kit (Biorad, 1725085) and Roche LightCycler 480 with PrimePCR SYBR green primer assays (Biorad) targeting the indicated transcripts: *Actb* (qMmuCED0027505), *Gapdh* (qMmuCED0027497), *Hprt* (qMmuCED0045738), *Stub1* (qMmuCED0001075), *Tbp* (qMmuCID0040542). Relative expression values were calculated using the ΔCT method correcting for PCR efficiency and mean centered across the three genotypes. Expression was normalized to the geometric mean of *Actb* and *Hprt*, the two most stable reference mRNAs across genotypes as determined via Normfinder (also tested: *Gapdh* and *Tbp*) [53].

#### Protein and mRNA isolation/analysis from mouse tissue

Intact tissues were isolated from anesthetized 4 month-old male littermates. Tissue was stored frozen in RNAlater solution (Ambion) until protein and mRNA were isolated using the Ambion PARIS Kit (Ambion, AM1921) as per manufacturer’s instructions. Tissue disruption prior to protein/RNA isolation was performed with Ambion PARIS Kit Cell Disruption Buffer and Qiagen TissueLyser LT with 5 mm steel beads. Any contaminating DNA was removed from RNA prepared by PARIS Kit by treatment with TURBO DNA-free™ Kit (Ambion) and mRNA was reverse transcribed using the iScript cDNA synthesis kit (Bio-Rad). Real-time PCR was performed using Roche LightCycler 480 and Sso Advanced Universal SYBR Green Supermix (Biorad) with PrimePCR SYBR green primer assays targeting the indicated mRNAs (Biorad) and methodology listed above. Expression was normalized to the geometric mean of *Actb* and *Gapdh*, the two most stable reference mRNAs across genotypes (also tested: *Hprt* and *Tbp*).

#### Cell lysate collection/nuclear fractionation/isolation of total, soluble and insoluble fractions

For all assays, unless otherwise noted, cell lysates were prepared by first washing cells in cold PBS and lysed in Cell Lytic M (Sigma) containing 1X HALT protease/phosphatase inhibitor (Pierce) and 50 µM PR619 (Lifesensors). Lysates were clarified by centrifugation at 15,000 x *g* for 10 min at 4 °C. Total protein concentration was determined by BCA protein assay (Pierce). Alternatively, cells were lysed on ice for 15 m in Triton X-100 cell lysis buffer (50 mM Tris, pH 7.4, 150 mM NaCl, 1% Triton X-100, 1mM EDTA, protease inhibitor (Complete; Roche), 50 µM PR-619 (LifeSensors). Triton X-100 insoluble material was collected by solubilization of the insoluble pellet following 15,000 x g centrifugation by resuspension in 2X Laemmeli Sample buffer (Biorad), brief sonication and heating for 5 m at 100 °C. Nuclear fractions were prepared using the NE-PER kit (Pierce), as per manufacturer’s instructions. Total, soluble and insoluble fractions as shown in were prepared by first trypsinizing and counting P2 primary MEFs grown on 15-cm tissue-culture treated dishes to near 100% confluency. Cells were then divided equally between two tubes, spun at 500 x *g*, pellet rinsed in PBS and spun again at 500 x *g*. Cells for total protein fraction (tube 1) were then lysed in 2X Laemmeli SDS sample buffer (65mM Tris-HCl, 10% Glycerol, 2% SDS), sonicated briefly on ice and boiled at 100 °C for 5 m. The cell pellet in tube 2 was then lysed for collection of soluble and insoluble fractions. This pellet was lysed in Triton X-100 cell lysis buffer as described above and soluble protein collected following centrifugation at 15,000 x *g* for 10 m at 4 °C. The insoluble pellet was then rinsed once in lysis buffer and spun again at 15,000 x *g* for 10 m at 4 °C. The pellet was then solubilized in 2X Laemmeli SDS sample buffer (65mM Tris-HCl, 10% Glycerol, 2% SDS), sonicated briefly on ice and boiled at 100 °C for 5 min. Total protein concentrations in each fraction were then determined by 660nm Protein Assay (Thermo Scientific) and samples of equal total protein prepared for SDS-PAGE by addition of final concentrations of 0.025% bromophenol blue and 100mM DTT.

#### Polyacrylamide gel electrophoresis, Blue native polyacrylamide gel electrophoresis, gel immunoblotting, and densitometry

For reduced and denatured conditions, protein samples were resolved on NuPAGE Novex® Bis-Tris Gels (Life Technologies) using the MOPS/LDS buffer system or Mini-PROTEAN® TGX Precast Gels (Bio-Rad) using the Tris/Glycine/SDS buffer. Native protein samples were resolved on 4-16% NativePAGE Novex® Gels (Life Technologies) using 0.001% G-250 cathode buffer. Proteins were transferred to PVDF membranes and incubated with primary antibodies overnight (see following table for antibody information) and detected with either anti-rabbit or anti-mouse (GE Healthcare), or anti-goat (Sigma) HRP-conjugated antibodies and visualized with ECL Advance substrate (GE Healthcare) using the EC3™ Imaging System (UVP). For quantification of relative protein levels, densitometry analysis was performed using LI-COR Image Studio Lite.

#### Immunoprecipitation/Cco-immunoprecipitation of AMPKα1/CHIP, Hsc70/CHIP from primary MEFs

Primary MEFs were cultured as described above and plated in 2 x 15 cm tissue culture dishes and incubated overnight under normal growth conditions. 24 h after plating, cells were treated with 20 µM MG132 or DMSO for 2.5 h prior to harvest. Cells were washed 1X in cold PBS and lysed in Cell Lytic M (Sigma) containing 1X HALT protease/phosphatase inhibitor (Pierce) and 50 µM PR619 (LifeSensors). Lysates were clarified by centrifugation at 15,000 x *g* for 10 min. Total protein concentration was determined by BCA protein assay (Pierce) and 1.5 mg total protein clarified lysates were incubated overnight at 4 °C with 10 µg of anti-AMPKα1 (R and D Systems, AF3197), anti-CHIP (Abcam, ab4447), rabbit IgG or goat IgG antibodies. 120 µl Protein-G Dynabeads (Invitrogen) was then added to each sample and incubated for 0.5 h at room temperature with rotation. Beads were washed four times with Phosphate-Buffered Saline with 0.05% Tween-20; subsequently, proteins were eluted in SDS-sample buffer and analyzed by SDS-PAGE and western blotting using anti-CHIP (Sigma, S1073), anti-AMPKα1/2 (Cell Signaling Technologies, 2532) or anti-Hsc70 (Enzo, ADI-SPA 815) antibodies.

#### CHIP immunofluorescence

Micrographs were obtained as previously described [54] with the following modifications. Cells were fixed in 4% paraformaldehyde for 10 min, then incubated in permeabilization buffer (PBS, 0.5% Triton X-100, 1% BSA) for 10 min. Primary and secondary antibodies were prepared at dilutions of 1:500 (CHIP-Sigma HPA043531) or (anti-c-myc, Sigma M4439) and 1:800 (Alexa-Fluor Goat anti-rabbit or Goat anti-mouse), respectively, in blocking buffer (PBS, 0.05% Triton X-100, 1% BSA). Coverslips were mounted using Vectashield Hardset with Dapi (Vector Laboratories). Cells were visualized using a Zeiss LSM 710 spectral confocal. Co-localization analysis was performed using the Coloc2 plugin (v 3.0.0) with Fiji [55, 56].

#### HSF1 luciferase reporter assay

COS-7 cells were cultured, plated at 5000 cells/well in clear-bottom white 96-well plates (Thermo Scientific, 165306) and transiently transfected as described above with the indicated transgene vectors and the Qiagen Cignal Heat Shock Response Reporter (luc) Kit (CCS4023L) as per manufacturer’s instructions. 24 h post-transfection cells, were lysed and luciferase assays were performed using a Dual-Luciferase® Reporter Assay System (Promega, Madison, WI) on a BMG Labtech CLARIOstar dual-injection plate reader following the manufacturer’s protocol. Transfection of each construct was performed in triplicate in each assay and a total of three assays were performed on three separate days. Empty vector was transfected in each plate in triplicate to be used for normalization purposes. Ratios of Renilla luciferase readings to Firefly luciferase readings were taken for each experiment and triplicates were averaged. The average values of the tested constructs were normalized to the activity of the empty construct.

#### Cycloheximide chase

COS-7 cells were co-transfected with the indicated vectors and 24 h post-transfection treated with 50 µg/ml cycloheximide for 0, 1 or 2.5 h in the presence or absence of 20 µM proteasome inhibitor MG132 and lysates collected and separated by SDS-PAGE and immunoblotted with antibodies against His-CHIP and β-tubulin as described above. Primary MEFs were plated and incubated for 24 h under normal growth conditions. Cells were then treated with for 0, 2, 4 or 6 h with 50 µg/ml cycloheximide and lysates collected and separated by SDS-PAGE and immunoblotted with antibodies against CHIP (ab4447) and β-tubulin as described above.

#### Heat shock and recovery

Two models of heat stress and recovery were utilized. For measures of heat stress/CHIP-induced nuclear translocation of HSF1 in COS-7 cells, cells were heat shocked in a water bath for 30 min at 42 °C prior to lysate collection and nuclear fractionation as previously described [27]. For Hsp70 induction and recovery assays in primary MEFs, cells were heat shocked in a water bath for 10 min at 42°C prior to recovery at 37°C under normal growth conditions for the indicated times.

### GENERATING CHIP-T246M PRECLINICAL MODELS AND BEHAVIORAL STUDIES

#### Data analysis

For all procedures, measures were taken by an observer blind to animal genotype. Behavioral data were analyzed using two-way repeated measures Analysis of Variance (ANOVA). If F values were < 0.05 on any individual factor or interaction of factors, Tukey’s multiple comparison test was used to compare means.

#### Generation of the equivalent CHIP-T246M at the endogenous rodent locus

The generation of the CHIP-T246M and CHIP-/-mice were previously described [27, 28]. CRISPR/Cas9 genetic editing was used to generate the same mutation in *Rattus norvegicus* (**Supplementary Figure S2A**).

##### Cas9-sgRNA targeting Plasmid construction

The precut pCS(puro) plasmid was used to express human codon-optimized Cas9 and sgRNA (**Supplementary Figure S2B**). The sgRNA were comprised of twenty nucleotides followed by the PAM sequence: Top oligo 5’-3’:CACCGGAACCCTGCATTACACCCAG; Bottom oligo 5’-3’ AAACCTGGGTGTAATGCAGGGTTCC. The precut pCS(puro) plasmid was ligated with the annealed oligos DNA to generate the Cas9-sgRNA targeting plasmids for STUB1 (**Supplementary Figure 2B**).

##### Donor vector construction

To minimize random integrations, we used a circular donor vector for homologous recombination repair. The targeting construct consisted of rat genomic fragment containing exons 1-6 of *Stub1* (**Supplementary Figure S2C**). The construct carried one nucleotide substitution in the initiation codon (ATG>ACA), and two homology arms of ∼2 kb each were used as templates to repair the double-strand breaks generated by Cas9. The donor vector was prepared using an endotoxin-free plasmid DNA kit.

##### Microinjection

Sprague Dawley (SD) female rats were used as embryo donors as well as for pseudo-pregnant foster mothers. Super-ovulated SD rats (3–4 weeks-of-age) were mated to SD males, and fertilized embryos were collected from the ampullae. Cas9-sgRNA targeting plasmids and donor vector were mixed at different concentrations and co-injected into the cytoplasm of fertilized eggs at the one-cell stage. After injection, surviving zygotes were transferred into the oviducts of SD pseudo-pregnant females.

##### Genotyping

Before weaning, all rats were genotyped by polymerase chain reaction (PCR) using tail DNA using a *Stub1* forward primer: 5’-CTCATGGGCAGGCTCTGGTATGG-3’; and a *Stub1* reverse primer 5’-GAGCAGTTCAGAACCCATCATCAGG-3’. PCR products were sequenced using sanger sequencing (**Supplementary Figure S2D**). All rats were genotyped confirmed for a second time, post-mortem.

#### Elevated plus maze for anxiety–like behavior

This procedure is based on a natural tendency of mice to actively explore a new environment, versus their fear of being in an open area. Mice were given one 5 min trial on the plus maze, which had two walled arms (the closed arms, 20 cm in height) and two open arms. The maze was elevated 50 cm from the floor, and the arms were 30 cm long. Animals were placed on the center section (8 cm x 8 cm), and allowed to freely explore the maze. Measures were taken of time on, and the number of entries into, the open and closed arms.

#### Marble-burying assay

Mice were tested in a Plexiglas cage located in a sound-attenuating chamber with ceiling light and fan. The cage contained 5 cm of corncob bedding, with 20 black glass marbles (14 mm diameter) arranged in an equidistant 5 x 4 grid on top of the bedding. Subjects were given access to the marbles for 30 min. Measures were taken of the number of buried marbles (two-thirds of the marble covered by the bedding).

#### Buried food test for olfactory function

Several days before the olfactory test, an unfamiliar food (Froot Loops, Kellogg Co., Battle Creek, MI) was placed overnight in the home cages of the mice. Observations of consumption were taken to ensure that the novel food was palatable. Sixteen to twenty hours before the test, all food was removed from the home cage. On the day of the test, each mouse was placed in a large, clean tub cage (46 cm L x 23.5 cm W x 20 cm H), containing paper chip bedding (3 cm deep), and allowed to explore for 5 min. The animal was removed from the cage, and 1 Froot Loop was buried in the cage bedding. The animal was then returned to the cage and given 15 min to locate the buried food. Measures were taken of latency to find the food reward.

#### Open field test

Exploratory activity in a novel environment was assessed by a one-hour trial in an open field chamber (41 cm x 41 cm x 30 cm) crossed by a grid of photobeams (VersaMax system, AccuScan Instruments). Counts were taken of the number of photobeams broken during the trial in 5-min intervals, with separate measures for ambulation (total distance traveled) and rearing movements. Time spent in the center region of the open field was measured as an index of anxiety-like behavior. Mice were tested at two ages: 8-9 weeks- and 32 weeks-of-age.

#### Rotarod

Mice and rats were tested for motor coordination and learning on an accelerating rotarod (Ugo Basile, Stoelting Co., Wood Dale, IL or Rot-Rod, Softmaze, respectively). For mice, the first test session animals consisted of three trials, with 45 s between each trial. Two additional trials were given 48 hours later. Rpm (revolutions per min) was set at an initial value of three, with a progressive increase to a maximum of 30 rpm across 5 min (the maximum trial length). Measures were taken for latency to fall from the top of the rotating barrel. Additional tests (two trials per test) were conducted at the indicated ages. For rats, each rat was placed on the Rotarod at a constant speed (5 rpm) for a maximum of 5 min, and at an accelerated speed (4 to 40 rpm in 5 min) for a maximum of 5 min. The latency to fall was recorded. Rats perform four trials for each time point, with 15 min rest between trials. For summary analysis, the mean latency to fall of each trial was used.

#### Sociability in a 3-chamber choice test

Mice were evaluated for differences in social preference. The test session consisted of three 10-min phases: a habituation period, a test for sociability, and a test for social novelty preference. For the sociability assay, mice were given a choice between being in the proximity of an unfamiliar conspecific (“stranger 1”), versus being alone. In the social novelty phase, mice were given a choice between the already-investigated stranger 1, versus a new, unfamiliar mouse (“stranger 2”). The social testing apparatus was a rectangular, 3-chambered box fabricated from clear Plexiglas. Dividing walls had doorways allowing access into each chamber. An automated image tracking system (Noldus Ethovision) provided measures of entries and duration in each side of the social test box, as well as time in spent within 5 cm of the Plexiglas cages (the cage proximity zone).

At the start of the test, the mouse was placed in the middle chamber and allowed to explore for 10 min, with the doorways into the two side chambers open. After the habituation period, the mouse was enclosed in the center compartment of the social test box, and an unfamiliar, sex-matched C57BL/6J mouse (stranger 1) was placed in one of the side chambers. The stranger mouse was enclosed in a small Plexiglas cage drilled with holes, which allowed nose contact, but prevented fighting. An identical empty Plexiglas cage was placed in the opposite side of the chamber. Following placement of the stranger and the empty cage, the doors were re-opened, and the subject was allowed to explore the entire social test box for 10 min. Measures were taken of the amount of time spent in each chamber and the number of entries into each chamber by the automated tracking system. At the end of the sociability phase, stranger 2 was placed in the empty Plexiglas container, and the test mouse was given an additional 10 min to explore the social test box.

#### Acoustic startle and prepulse inhibition

This procedure was used to assess auditory function, reactivity to environmental stimuli, and sensorimotor gating. The test was based on the reflexive whole-body flinch, or startle response, which follows exposure to a sudden noise. Measures were taken of startle magnitude and prepulse inhibition, which occurs when a weak prestimulus leads to a reduced startle in response to a subsequent louder noise. Mice were tested at two ages: 11-13 weeks and 33 weeks. For each test, mice were placed into individual small Plexiglas cylinders within larger, sound-attenuating chambers. Each cylinder was seated upon a piezoelectric transducer, which allowed vibrations to be quantified and displayed on a computer. The chambers included a ceiling light, fan, and a loudspeaker for the acoustic stimuli. Background sound levels (70 dB) and calibration of the acoustic stimuli were confirmed with a digital sound level meter (San Diego Instruments). Each session consisted of 42 trials that began with a 5-min habituation period. There were seven different types of trials: the no-stimulus trials, trials with the acoustic startle stimulus (40 msec; 120 dB) alone, and trials in which a prepulse stimulus (20 msec; either 74, 78, 82, 86, or 90 dB) occurred 100 ms before the onset of the startle stimulus. Measures were taken of the startle amplitude for each trial across a 65-msec sampling window, and an overall analysis was performed for each subject’s data for levels of prepulse inhibition at each prepulse sound level (calculated as 100 - [(response amplitude for prepulse stimulus and startle stimulus together / response amplitude for startle stimulus alone) x 100].

#### Fear conditioning

Mice were evaluated for learning and memory in a conditioned fear test, using the Near-Infrared image tracking system (MED Associates, Burlington, VT). The procedure had the following phases: training on Day 1, a test for context-dependent learning on Day 2, and a test for cue-dependent learning on Day 3. Follow-up tests for retention of learning were conducted 2 weeks later.

##### Training

On Day 1, each mouse was placed in the test chamber, contained in a sound-attenuating box, and allowed to explore for 2 min. The mice were then exposed to a 30 s tone (80 dB), followed by a 2 s scrambled foot shock (0.4 mA). Mice received two additional shock-tone pairings, with 80 s between each pairing.

##### Context- and cue-dependent learning

On Day 2, mice were placed back into the original conditioning chamber for a test of contextual learning. Levels of freezing (immobility) were determined across a 5 min session. On Day 3, mice were evaluated for associative learning to the auditory cue in another 5 min session. The conditioning chambers were modified using a Plexiglas insert to change the wall and floor surface, and a novel odor (dilute vanilla flavoring) was added to the sound-attenuating box. Mice were placed in the modified chamber and allowed to explore. After 2 min, the acoustic stimulus (an 80 dB tone) was presented for a 3 min period. Levels of freezing before and during the stimulus were obtained by the image tracking system.

Two weeks following each test, mice were given second tests to evaluate retention of context- and cue-dependent learning.

#### Gait analysis

The CatWalk XT (Noldus information Technology, Wageningen, Netherlands) was used to analyze gait of unforced moving rats. CatWalk XT consists a hardware system of a long glass walkway plate, illuminated with green light that is reflected within the glass at points be touched, a high-speed video camera, and a software package for quantitative assessment of animal footprints. The parameters we observed included stride length: the distance between successive placements of the same paw; base of support: the average width between either the front paws or the hind paw; step cycle: the time in seconds between two consecutive initial contact of the same paw. All rats were trained to cross the runway in consistently at least six times a day for a week before any experimentation. A successful run was defined as an animal finishing the run down the tracks without any interruption or hesitation. Rats that failed the CatWalk training were excluded from the study. An average number of five replicate crossings by each rat was recorded. Rats were subjected to computer-assisted CatWalk monthly after 8 weeks-of-age.

#### Morris water maze

The Morris water maze task was used to assess learning and memory. The task was conducted in a round tank, 160 cm in diameter and 54 cm deep, filled with water. The wall was colored with non-toxic black paint to ensure opaqueness. Throughout testing, the water temperature was monitored and maintained at 21 ℃. The tank was divided into four equally sized quadrants, and a circular acrylic escape platform was placed in one of the quadrants. The escape platform was submerged in water by 2 cm so that it was not visible to the animals. A camera mounted above the tank recorded the movement of the animals in each trial. The Sunny Instruments Morris water maze Tracking Software was used to record the latency to reach the escape platform and the time spent in the target quadrant. The water maze task consisted of four training days with four trials on each day. In each trial, the animals were placed in the water facing the tank wall and had to locate the escape platform. The initial position of the animal was the vertices of one of the four quadrants and was different for each trial. It was assigned randomly and counterbalanced for the genotypes. Animals could utilize external visual cues on the walls surrounding the tank to locate the platform. The trial was completed when the animal either found the escape platform or 60 s had passed. If the animal was unable to locate the platform in 60 s, it was gently led to the platform. Animals were allowed to remain on the escape platform for 15 s before being removed and dried for the next trial. In addition, the platform position was kept constant between trials and days. The four trials of the first four training days were used as an indicator of spatial working memory. On day 5, the animals performed a 60 s probe trial without the platform. During this trial, the time spent in the target quadrant was recorded for each animal.

#### Study approval

All animal procedures were approved by the Institutional Animal Care and Use Committee of The University of North Carolina at Chapel Hill or the Medical Research Ethics Committee Board of the First Affiliated Hospital of Zhengzhou University.

### PROTEOMIC STUDIES

#### Sample preparation, iTRAQ labeling, and LC-MS/MS analysis

Immediately after termination, the cerebellum was dissected, briefly washed with PBS solution, snap-frozen in liquid nitrogen, and stored at -80 °C until use. Frozen samples were obtained from both genotypes (N = 3) and total proteins of each sample were extracted using lysis buffer (1% SDS, 50 mM Tris–HCl (pH 6.8), 10% glycerol, and 1× protease inhibitor). Lysates were boiled in water for 15 min, sonicated (80 W, pulse at 10 s then 15 s for 10 times), and then boiled again for 15 min. Finally, lysates were centrifuged at high speed for 30 min, and supernatants collected and stored in aliquots at --80 °C (longer term)

Protein solutions (100 µg) containing 8 M urea was diluted four times with 100 mM Triethylammonium bicarbonate. Trypsin Gold (Promega, Madison, WI, USA) was used to digest the proteins with the ratio of protein: trypsin =40:1 at 37 °C overnight. After trypsin digestion, peptides were desalted with a Strata X C18 column (Phenomenex) and vacuum-dried according to the manufacturer’s protocol. Samples were labeled using the iTRAQ Reagent-8plex Multiplex Kit (AB SCIEX, Framingham, USA), according to the manufacturer’s instructions. T246/T246 samples were labeled with iTRAQ tags 117, 118, and 119, while M246/M246 samples were labeled with tags 114, 115, and 116 (three replicates from each group). After 2 h of incubation at room temperature, labeled samples were mixed at equal ratios, desalted with a Strata X C18 column (Phenomenex), and vacuum-dried according to the manufacturer’s protocol.

The peptides were separated on a Shimadzu LC-20AB HPLC Pump system coupled with a high pH RP column. The peptides were reconstituted with buffer A (5% acetonitrile, 95% H2O, adjusted pH to 9.8 with ammonia) to 2 ml and loaded onto a column containing 5-µm particles (Phenomenex). The peptides are separated at a flow rate of 1 mL/min with a gradient of 5% buffer B (5% H2O, 95% acetonitrile, adjusted pH to 9.8 with ammonia) for 10 min, 5-35% buffer B for 40min, 35-95% buffer B for 1 min. The system was then maintained in 95% buffer B for 3 min and decreased to 5% within 1 min before equilibrating with 5% buffer B for 10 min. Elution is monitored by measuring absorbance at 214 nm, and fractions are collected every 1 min. The eluted peptides are pooled as 20 fractions and vacuum dried.

Each fraction was resuspended in buffer A (2% acetonitrile and 0.1% formic acid in water) and centrifuged at 20,000 g for 10 min. The supernatant was loaded onto a C18 trap column 5μl/min for 8 min using an LC-20AD nano-HPLC instrument (Shimadzu, Kyoto, Japan) by the autosampler. Then, the peptides were eluted from trap column and separated by an analytical C18 column (inner diameter 75 µm) packed in-house. The gradient was run at 300 nl/min starting from 8 to 35% of buffer B (2% H2O and 0.1% formic acid in acetonitrile) in 35 min, then going up to 60% in 5 min, then maintenance at 80% B for 5 min, and finally return to 5% in 0.1 min and equilibrated for 10 min.

Data acquisition was performed with a TripleTOF 5600 System (SCIEX, Framingham, MA, USA) equipped with a Nanospray III source (SCIEX, Framingham, MA, USA), a pulled quartz tip as the emitter (New Objectives, Woburn, MA) and controlled with software Analyst 1.6 (AB SCIEX, Concord, ON). Data were acquired with the following MS conditions: ion spray voltage 2,300 V, curtain gas of 30, nebulizer gas of 15, and interface heater temperature of 150 °C. High sensitivity mode was used for the whole data acquisition. The accumulation time for MS1 is 250ms, and the mass ranges were from 350 to 1500 Da. Based on the intensity in MS1 survey, as many as 30 product ion scans were collected exceeding a threshold of 120 counts per second (counts/s) and with charge-state 2+ to 5+. The dynamic exclusion was set for 1/2 of peak width (12 s). For ITRAQ data acquisition, the collision energy was adjusted to all precursor ions for collision-induced dissociation, and the Q2 transmission window for 100Da was 100%.

#### Protein identification

The raw MS/MS data were converted into MGF format by ProteoWizard tool msConvert, and the exported MGF files were searched using Mascotversion 2.3.02 in this project against the selected database. At least one unique peptide was necessary for the identified protein. The databases used were: NCBInr, SwissProt, UniProt. An automated software called IQuant was used for quantitatively analyzing the labeled peptides with isobaric tags. It integrates Mascot Percolator, a well-performing machine learning method for rescoring database search results, to provide reliable significance measures. To assess the confidence of peptides, the PSMs were pre-filtered at a PSM-level FDR of 1%. Then based on the “simple principle” (the parsimony principle), identified peptide sequences are assembled into a set of confident proteins. To control the rate of false-positive identifications at the protein level, an FDR threshold of 1% was used, which is based on Picked protein FDR strategy, will also be estimated after protein inference (protein-level FDR <= 0.01). The protein quantification process includes the following steps: protein identification, tag impurity correction, data normalization, missing value imputation, protein ratio calculation, statistical analysis, results presentation. Protein identification was supported by all peptide matches with 95% confidence. Comparisons were made between genotypes and proteins were considered to be differentially that met the criteria of *p* < 0.05 and absolute fold change > 1.2.

#### StringDB analysis

Differentially expressed proteins were analyzed using STRING analysis [37]. We filter excluded any proteins lacking interactions and used the Markov cluster algorithm (MCL) with an inflation parameter of three to identify clusters of related proteins based on their interaction network. Simultaneously, we used functional enrichment to identify biological processes, cellular components, and pathways that were over-represented in our protein list using a false discovery rate of less than 5%.

## Acknowledgments

We thank members of the Schisler Laboratory for critical review of the manuscript; the Willis, Jensen, McLean, and Stouffer Laboratories for support; and The McAllister Heart Institute administration team. We thank John B. Belcher, Sabrina Madrigal, Saranya Ravi, Alex Eaker, and Anna Beth Robertson for their help with animal studies. We also thank the core facilities at The University of North Carolina at Chapel Hill for their services, including the Mouse Behavioral Phenotyping Core (Sheryl Moy, PhD), the Animal Histopathology and Lab Medicine Core (Dawud Hilliard), and the Macromolecular Interactions Facility (Ashutosh Tripathy, PhD). This study was funded by grants 81530037 and 81471158 (YX), and 81771290 (CS) from the National Natural Science Foundation of China; and R37 HL065619 from the National Institutes of Health (to CP and JCS). All authors approved the final version of the manuscript and agree to be accountable for all aspects of the work in ensuring that questions related to the accuracy or integrity of any part of the work are appropriately investigated and resolved. All persons designated as authors qualify for authorship, and all those who qualify for authorship are listed.

## Video legends

**Video 1:** *Performance on the accelerating rotoarod is decreased in adult rats homozygous for CHIP-T246M.* Three rats with either the M246/M246 (left) or T246/T246 (right) genotypes were placed on the non-moving rod. Once the rod starts to rotate, M246/M246 mice fall off within 15 s.

**Video 2:** *Hind limb clasping in mice with disrupted CHIP expression.* (**A**) Mice with the M246/M246 (left, blue glove) or T246/T246 (right, white glove) genotype or (**B**) mice that lack CHIP expression (left, blue glove) or a wild-type litter mate (right, white glove), were tested for the degree of hind limb clasping. Increased clasping was observed in M246/M246 and CHIP-/- mice and further increased with age.

## Supplementary Tables

**Supplementary Table S1:** *Proteomics analysis of differentially expressed cerebellar proteins identified in the CHIP-T246M rat model.* The protein name is provided along with the values of each protein from biological replicates of M246/M246 cerebellums (M1, M2, and M3) relative to the mean values of three T246/T246 biological replicates. The mean of all three replicates, the standard deviation (SD), and the *p* value comparing M246/M246 vs. T246/T246 conditions are indicated. Additional annotations of each protein, including database ID, the mass, and decryption are provided.

**Supplementary Table S2:** Cluster analysis of interacting proteins. Each cluster was color coded as found in Figure 7B. The number of proteins in each cluster, the protein name, ID, and description is provided.

## Supplementary Figures

**Supplementary Figure S1:**
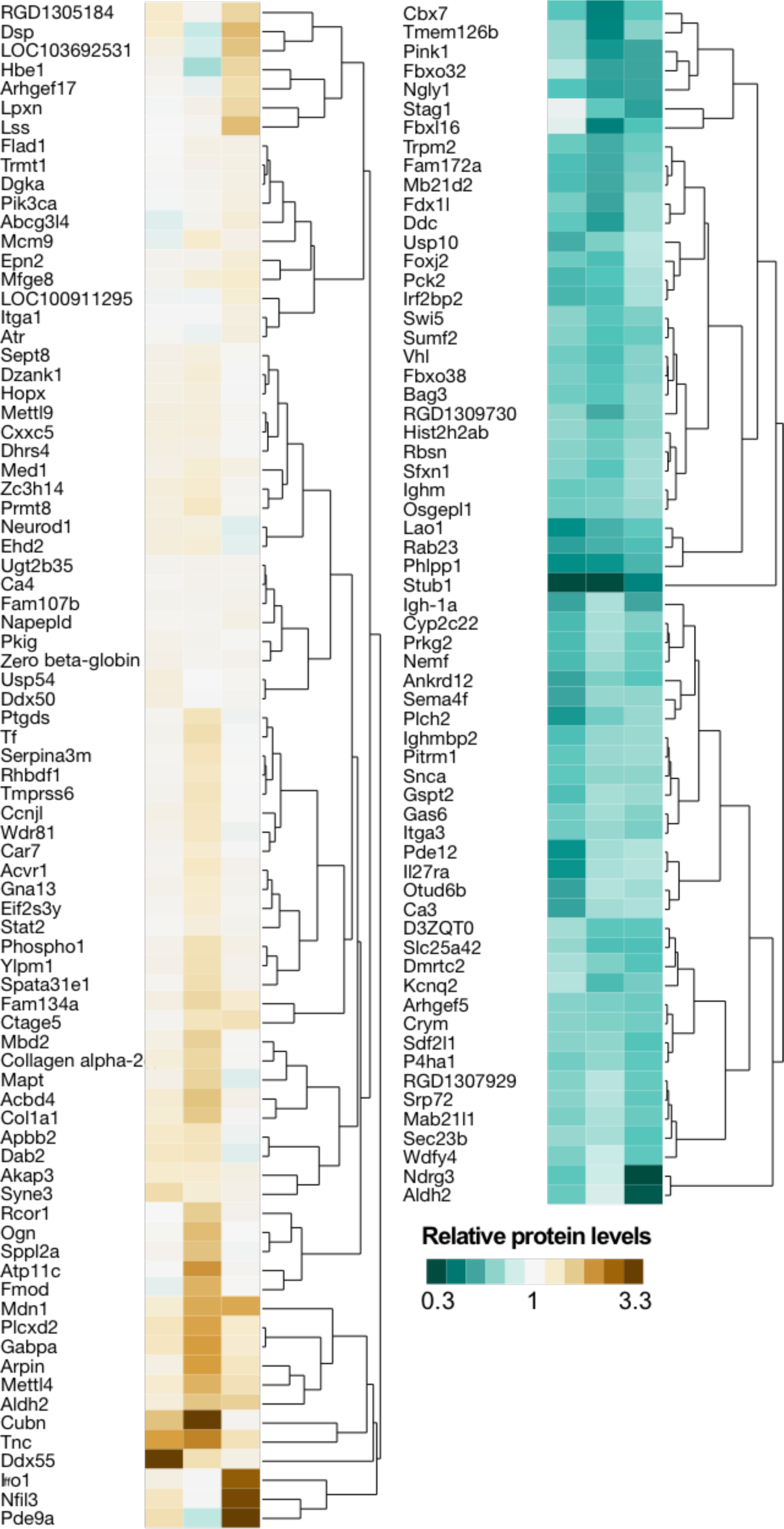
Unsupervised cluster analysis of differentially expressed proteins due to CHIP-T246M. Proteins were clustered using Ward linkage analysis. Each column is the mean of a biological replicate of an M246/M246 cerebellum relative to T246/T246 control cerebellums. The two primary clusters represent proteins that are either increased (left) or decreased (right) in M246/M246 cerebellums as indicated by the color bar.

**Supplementary Figure S2:**
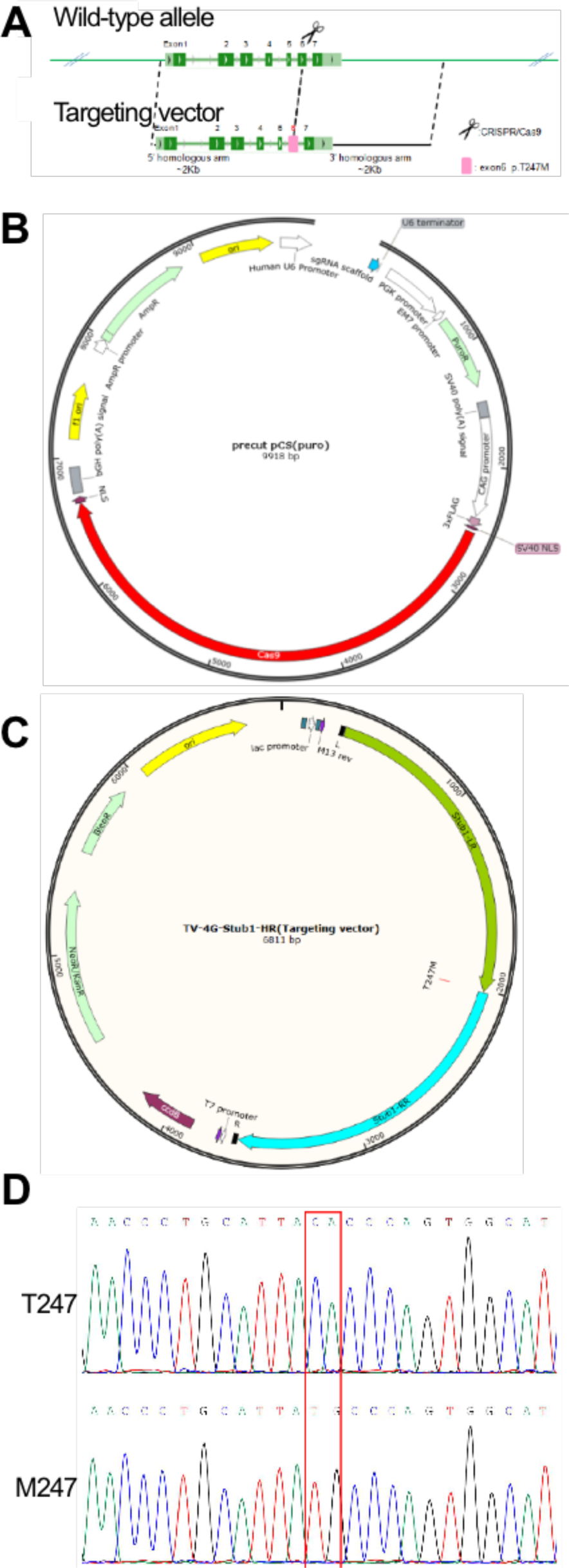
Generation the corresponding CHIP-T246M mutation at the endogenous locus of Stub1 in Rattus norvegicus using CRISPR/Cas9. (A) Schematic of targeting vector used for in vivo genome editing, targeting the Cas9 nuclease to exon 6. (**B**) Map of Cas9 vector. (**C**) Map of *Stub1* T247M targeting vector (rodents have an additional coding exon relative to humans). (**D**) Sanger sequencing confirmation of the T246M mutation, changing the coding exon from ACA (threonine) to ATG (methionine).

